# What’s past is past, mostly: *Brassicaceae* host plants mask the feedback from the previous year’s soil history on bacterial communities, except when the *Brassicaceae* hosts experience drought

**DOI:** 10.1101/2021.12.07.471648

**Authors:** Andrew J.C. Blakney, Luke D. Bainard, Marc St-Arnaud, Mohamed Hijri

**Affiliations:** Institut de recherche en biologie végétale, Université de Montréal and Jardin botanique de Montréal, Montréal, QC, Canada; Agassiz Research and Development Centre, AgricuSlture and Agri-Food Canada, Agassiz, BC, Canada; African Genome Center, Mohammed VI Polytechnic University (UM6P), Lot 660, Hay Moulay Rachid, Ben Guerir 43150, Morocco

**Keywords:** *Brassicaceae*, bacterial communities, soil legacy, plant-soil feedbacks

## Abstract

Previous soil history and the current plant hosts are two plant-soil feedbacks that operate at different time-scales to influence the structure soil bacterial communities. In this study, we used a MiSeq metabarcoding strategy to describe the impact of five *Brassicaceae* host plant species, and three different soil histories, on the structure of their bacterial root and rhizosphere communities at full flower. We found that the *Brassicaceae* host plants were consistently significant in structuring the bacterial communities. Four host plants (*Sinapis alba, Brassica napus, B. juncea, B. carinata*) formed nearly the same bacterial communities, regardless of soil history. *Camelina sativa* host plants structured phylogenetically distinct bacterial communities compared to the other hosts, particularly in their roots. Soil history established the previous year was only a significant factor for bacterial community structure when the feedback of the *Brassicaceae* host plants was weakened, potentially due to limited soil moisture during a dry year. Understanding how plant-soil feedbacks operate at different time-scales and are involved in how microbial communities are structured is a pre-requisite for employing microbiome technologies in improving agricultural systems.

## Introduction

Previous soil history and the current plant hosts are two aspects involved in structuring soil microbial communities (Fitzpatrick *et al*., 2018; Hannula *et al*., 2021). Microbial communities manage the ecological function of the soil, and its biological stability (Griffiths & Philippot, 2013). Soil microbes, bacteria in particular, aggregate, structure, and stabilize soils (Duchicela et al., 2013; Graf & Frei, 2013), as well as water and nutrient cycling through the biosphere (Talbot et al., 2013). In relation to plants, soil bacteria increase access to nutrients (Richardson et al., 2009), temper environmental change (Lau & Lennon, 2012), or stress (Marasco et al., 2012), and protect against pathogens (Mendes et al., 2011; Sikes et al., 2009). Therefore, in order to improve how plants interact with their bacterial communities, we must understand the rules that structure bacterial communities. Without this knowledge, plant microbial communities cannot be predicted, design, or deployed for future needs (Albright *et al*., 2021, Busby *et al*., 2017, Zengler *et al*., 2019). Ultimately, understanding how plants interact with their bacterial communities are paramount for maintaining global biodiversity, agricultural productivity, and climate management (Albright *et al*., 2021). As such, a clear barrier to successfully using bacterial technologies, is proper understanding of the role of soil history and the individual plant hosts in environmental filtering (Albright *et al*., 2021, Busby *et al*., 2017).

How specific bacterial taxa come to inhabit a given site is an on-going process? The environmental and chemical context of the soil is the first step in establishing an environmental filter, which will begin to select for certain bacteria. Plants that subsequently grow in this site will also modify the environmental filter, as plant growth, development, and homeostasis, is determined by a plant’s capacity to uptake nutrients from the soil, altering the soil chemistry (Hu *et al*., 2021). Healthy plants will continue to modify the local soil by rhizodeposition (Kawasaki *et al*., 2021). This process of establishing an environmental filter is dynamic through time, as demonstrated by plant-soil feedbacks (PSFs), therefore, the bacterial community of a “current” resident plant can be influenced by a “previous” plant, or event, such as a drought, or fertilization (Kaisermann *et al*., 2017).

Agricultural systems provide a ready-made opportunity to investigate how PSFs impact bacterial community ecology. For example, crop rotations demonstrate how a previous plant impacts a current plant; introducing legumes, like lentils, into a rotation, tends to give higher yields for the subsequent plant, possibly due to increasing soil moisture, and/or available nitrogen (O’Donovon *et al*., 2014; Hamel *et al*., 2018). Crop rotations establish PSFs, and though extensive work has investigated how they benefit the plants involved (recently reviewed by Yang *et al*., 2021), how PSFs in crop rotations, and their agricultural inputs, impact soil microbial communities has been less examined.

Recent work on the canola (*Brassica napus* L. or *B. juncea* L.) bacterial community has begun to correct this knowledge deficit (Lay *et al*., 2018; Floch *et al*., 2020; Taye *et al*., 2020; Wang *et al*., 2020; Morales Moreira *et al*., 2021). *Brassicaceae* oilseed-based rotations are common throughout the world, as demand for vegetable oil and biofuels increase (Yang *et al*., 2021). Expanding the diversity of *Brassicaceae* oilseed species has been on-going, in order to improve production by identifying plants resistant to pathogens, and better adapted to the heat and drought stress of the Canadian Prairies (Bailey-Serres *et al*., 2019; Hossain *et al*., 2019; Liu *et al*., 2019). Studying closely related *Brassicaceae* species will help illustrate how bacterial communities may be changed through plant breeding, as well as the fine-tuning of bacterial community structures in response to PSFs, between two growing seasons, as well as within one growing season (Bailey-Serres *et al*., 2019).

To study the impact of PSFs at different time-scales, we took advantage of an existing agriculture experiment to investigate the impact of the previous year’s soil history, and the current host plants, on soil bacterial communities Three soil histories were established by growing wheat, lentils, or being left fallow (Fig. S1A). The following year, each soil history was divided into five subplots, and planted with a different *Brassicaceae* oilseed species. At full flower, which corresponds to the high activity of rhizosphere microbial communities, the root systems of these *Brassicaceae* host plants were harvested, and divided into root and rhizosphere compartments, from which the environmental DNA was extracted. We used a MiSeq metabarcoding approach to sequence the 16S rRNA gene from the bacterial communities for each root and rhizosphere sample, where the sequencing data was used to infer amplicon sequence variants (ASVs) to identify the bacterial biodiversity (Fig. S2).

The influence of PSFs is expected to fade through time, where the current host plant ought to determine the structure of the bacterial community (Hannula *et al*., 2021) Here, we investigated how PSFs at two different time-scales, the soil history established the previous year, and the current year’s host plants, vary in their influence on structuring the bacterial rhizosphere and root communities. Specifically, we tested two key hypotheses: 1) that each of the five current *Brassicaceae* host plants would structure unique bacterial communities in their roots and rhizosphere, and 2) that the three soil histories established the previous year would impact the structure of these bacterial communities differently. To test these hypotheses we examined the taxonomic composition and diversity of bacterial communities from the root and rhizosphere, as estimated by amplicon sequence variants (ASVs), to identify any changes associated with soil history, or the *Brassicaceae* hosts. Our results demonstrated that the influence of the current *Brassicaceae* host plants masked the influence on the previous year’s soil history in structuring the bacterial communities. When the influence of the current *Brassicaceae* host plants was weakened, then the impact from the previous year’s soil history could be detected, with some differences between communities with different soil histories. Moreover, we found that only *Camelina sativa* host plants established phylogenetically distinct bacterial communities, compared to the four other *Brassicaceae* host plants (*Sinapis alba, Brassica napus, B. juncea, B. carinata*) that formed similar bacterial communities, regardless of soil history.

## Materials & Methods

### Site and experimental design

A field experiment was conducted at the experimental farm of Agriculture and Agri-Food Canada’s Research and Development Centre, in Swift Current, Saskatchewan (50°15′N, 107°43′W). The site is located in the semi-arid region of the Canadian Prairies; according to the weather station of the research farm, the 2016 and 2017 growing seasons (May, June and July) had 328.4 mm and 55.0 mm of precipitation, respectively; compared to the 30-year average [1981-2010] of 169.2 mm. The daily temperature averages for the 2016 and 2017 seasons were 15.6°C and 15.9°C, respectively, while the 30-year average was 14.93°C. The farm is on a Brown Chernozem with a silty loam texture (46% sand, 32% silt, and 22% clay), and has been well-described previously (see Liu *et al*., 2019 & 2020).

The experiment was established in a field previously growing spring wheat (*Triticum aestivum* cultivar AC Lillian). A two-year (or two-phase) cropping sequence was repeated twice, Experiment 1, 2015-2016, and Experiment 2, 2016-2017, on adjacent sites (Fig. S1B & C). On each site, the experimental design was a split-plot replicated in four complete blocks. In the ‘Conditioning’ Phase, three soil history treatments were randomly assigned to the main plots, consisting of spring wheat (*Triticum aestivum*, cv. AC Lillian), red lentil (*Lens culinaris* cv. CDC Maxim CL), or left fallow (Fig. S1A). In Phase 2, the ‘Test’ Phase, the 12 Conditioning Phase plots were each subdivided and five *Brassicaceae* oilseed crop species were randomly assigned to one of these five subplots (Fig. S1A & S1B). The *Brassicaceae* crops seeded were Ethiopian mustard (*Brassica carinata* L., cv. ACC110), canola (*B. napus* L., cv. L252LL), oriental mustard (*B. juncea* L., cv. Cutlass), yellow mustard (*Sinapis alba* L., cv. Andante), and camelia (*Camelina sativa* L., cv. Midas). In total, each experiment had 60 subplots to sample (Fig. S1B). For further details of this well-described experiment, its design, and treatments, see Hossain *et al*. (2019), Liu *et al*. (2019), and Wang *et al*. (2020).

### Crop management and sampling

Conditioning and Test Phase crops were grown and maintained according to standard management practices, as previously described by Hossain *et al*. (2019), Liu *et al*. (2019), and Wang *et al*. (2020). A pre-seed ‘burn off’ herbicide treatment using glyphosate (Roundup, 900 g acid equivalent per hectare, a. e. ha^−1^) was applied to all plots each year to ensure a clean starting field prior to seeding. Conditioning Phase lentil seeds were treated with a commercial rhizobium-based inoculant (TagTeam at 3.7 kg ha^−1^). Conditioning Phase lentil and wheat were direct-seeded into wheat stubble from late April to mid-May, depending on the crop and year. The herbicides, glyphosate (Roundup, 900 g a. e. ha^−1^), Assure II (36 g active ingredient per hectare, a. i. ha^−1^), and Buctril M (560 g a.i. ha^−1^) were applied to the fallow, lentil, and wheat plots, respectively, for in-season weed control, while fungicides were only applied as needed. Soil tests were used to determine the rates of in-season nitrogen, phosphorus, and potassium, application; no synthetic nitrogen fertilizer was applied to the lentil plots (Table S4). Both lentil and wheat were harvested between late August and early October, depending on the crop and year. The Conditioning Phase established a soil history composed of either wheat, lentil, or fallow, plus their respective management plans (Hossain *et al*., 2019; Liu *et al*., 2019).

The subsequent Test Phase *Brassicaceae* plant hosts were subjected to the same standard management practices as the Conditioning Phase, including pre-seed ‘burn off’, in-season herbicide and fungicide treatments as needed, and fertilized as recommended by soil tests (Table S4; Hossain *et al*. 2019, Liu *et al*. 2019, and Wang *et al*. 2020). Additionally, all *Brassicaceae* crops, except *B. napus*, were treated with Assure II mixed with Sure-Mix or Merge surfactant (0.5% v/v) for post-emergence grass control: Liberty (glufosinate, 593 g a.i. ha−1) was used for *B. napus*. The Test Phase established the *Brassicaceae* host effect, composed of the individual *Brassicaceae* genotypes, as well their respective management plans.

Test phase *Brassicaceae* crops were sampled in mid-late July at full flowering, i.e. when 50% of the flowers on the main raceme were in bloom, as described by the Canola Council of Canada (Canola Encyclopedia: Canola Growth Stages, 2017). Four plants from two different locations within each subplot were excavated and pooled together as a composite sample (Hossain *et al*., 2019; Liu *et al*., 2019, Wang *et al*., 2020). In the field, each plant had its rhizosphere soil divided from the root material by gently scraping it off using bleach sterilized utensils into fresh collection trays. The roots were then gently washed three times with sterilized distilled water to remove any soil. Both the rhizosphere and root portions were immediately flash-frozen and stored in liquid nitrogen vapour shipping containers until stored in the lab at −80°C (Delavaux *et al*., 2020). Based on the sampling strategy, in this study we define the rhizosphere microbiome as the microbial community in the soil in close contact with the roots, and the root microbiome as the microbial community attached to, and within, the roots. Additional soil material was collected from each plot, kept on ice in coolers, and homogenized in the lab. These soil samples were used for soil chemistry analyses, including total carbon, nitrogen, pH, and micronutrients (see Wang *et al*., 2020 for details). Aerial portions of each harvested plant sample were retained to determine dry weight (Fig. S3).

### DNA extraction from Test Phase *Brassicaceae* root and rhizosphere samples

Nucleic acids were extracted from Experiment 1 Test Phase *Brassicaceae* samples, both rhizosphere and root portions. First, all the root samples were ground in liquid nitrogen via sterile mortar and pestles (Fig. S2). Total DNA and RNA were extracted from ∼1.5 g of rhizosphere soil using the RNA PowerSoil Kit with the DNA elution kit (Qiagen, Germany). DNA and RNA were extracted using ∼0.03 g of roots using the DNeasy Plant DNA Extraction Kit, and RNeasy Plant Mini Kit (Qiagen, Canada), respectively, following the manufacturer’s instructions (see Wang *et al*. (2020) for use of the RNA samples). All remaining harvested material from Experiment 1 and 2 Test Phase, as well as the extracted DNA from Experiment 1 Test Phase samples, were kept at − 80°C before being shipped to Université de Montréal’s Biodiversity Centre, Montréal (QC, Canada) on dry ice for further processing (Delavaux *et al*., 2020; Lay *et al*., 2018).

Total DNA was extracted from the Experiment 2 Test Phase samples; ∼500 mg of rhizosphere soil was used for the NucleoSpin Soil gDNA Extraction Kit (Macherey-Nagel, Germany), and ∼130 mg of roots for the DNeasy Plant DNA Extraction Kit (Qiagen, Germany) (Lay *et al*., 2018). A no-template extraction negative control was used with both the root and rhizosphere extractions and included with the Test Phase samples (Fig. S2), to assess the influence of the extraction kits on our sequencing results, and the efficacy of our lab preparation. All 242 extracted DNA samples (60 plots x 2 parallel experiments x 2 compartments, rhizosphere and root, +2 no-template extraction control samples) were quantified using the Qubit dsDNA High Sensitivity Kit (Invitrogen, USA), and qualitatively evaluated by mixing ∼2 μL of each sample with 1 μL of GelRed (Biotium), and running it on a 0.7 % agarose gel for 30 minutes at 150 V. The no-template extraction negative controls were confirmed to not contain detectable amounts of DNA after extraction. Samples were kept at −80°C (Bell *et al*., 2016; Delavaux *et al*., 2020).

### 16S rRNA gene amplicon generation and sequencing to estimate the bacterial community

To estimate the composition of the bacterial communities in the rhizosphere and roots from the Test Phase *Brassicaceae* species, extracted DNA from all samples were used to prepare 16S rRNA gene amplicon libraries following Illumina’s MiSeq protocols. First, all DNA samples were diluted 1:10 into 96-well plates using the Freedom EVO100 robot (Tecan, Switzerland). To assess potential bias caused by lab manipulations, sequencing and downstream bioinformatic processing, a commercially available 16S rRNA mock community, of known composition (Table S1), was included on each plate (Fig. S2) following the manufacturer’s instructions (BEI Resources, USA). The mock community contained DNA of 20 bacterial species (Table S2) in equimolar counts (10^6^ copies/μL) of 16S rRNA genes.

DNA plates were stored at −20°C before 2 μL from each sample was used as template in the 16S rRNA PCR reactions (see Supplementary Methods for details), set-up in 96-well plates using the Freedom EVO100 robot (Tecan, Switzerland). A no-template PCR negative control was included on each plate, to assess the influence of the PCR reaction, and the efficacy of our lab preparation on sequencing (Fig. S2). Each sample, and all controls, were PCR amplified in two independent reactions, except four rhizosphere samples from 2017, which we were unable to amplify, and were subsequently excluded hereafter (Fig. S2). Four μL of each reaction product was mixed with 1 μL of loading dye containing Gel Red (Biotium), and visualized on a 1% agarose gel after 60 minutes at 100 V. None of the no-template negative controls, from either the extractions, or the PCR reactions, contained detectable amounts of DNA after PCR amplification. All samples were then cleaned using the NucleoMag NGS Clean-Up Kit (Macherey-Nagel, Germany), following the manufacturer’s instructions. The cleaned products of the duplicated 16S rRNA gene PCR reaction, were then pooled together and submitted for paired-end 250 bp sequencing using Illumina’s MiSeq platform (Genome Québec, Montréal) (Bell *et al*., 2016; Lay *et al*., 2018). We estimated this should provide a mean of 39 000 reads per sample, which is in excess of our previous work to describe bacterial communities (Bell *et al*., 2016; Lay *et al*., 2018).

### Estimating ASVs from MiSeq 16S rRNA gene amplicons

The 16S rRNA gene amplicons generated by Illumina MiSeq were used to estimate the diversity and composition of the bacterial communities present in both the rhizosphere and root of each Test Phase *Brassicaceae* sample. The integrity and totality of the 16S MiSeq data downloaded from Génome Québec was confirmed using their MD5 checksum protocol (Roy *et al*., 2018). Subsequently, all data was managed, and analyzed in R (4.0.3 R Core Team, 2020), and plotted using `ggplot2` (Wickham, 2016).

Instead of generating OTUs from the 16S rRNA gene amplicon data, we opted to use DADA2 for ASV inference, as it generates fewer false-positives than OTUs, and reveals more low-abundant, or cryptic, microbes. Moreover, as ASVs are unique sequence identifiers, they are directly comparable between studies, unlike OTUs (Callahan *et al*., 2016a & 2017; Fitzpatrick *et al*., 2018). The `dada2` package (Callahan *et al*., 2016a) was first used to filter and trim all 23 313 756 raw reads, forward and reverse, from the 16S rRNA gene amplicon data generated from the control samples, the mock communities, and the Test Phase *Brassicaceae* samples, using the `filterAndTrim` function (Fig. S2). All reads were trimmed from the 3’ end to 240 bp, as determined *a posteriori* to be the optimum length to maximize retention of reads (14 666 924 after filtration, 10 178 467 retained overall), and the subsequent inference of ASVs (37 445). Any residual sequencing artifacts, including primers, were trimmed from the 5’, and low-quality (Q = < 20) reads removed, before ASVs were inferred (Fig. S4C).

Filtered and trimmed reads were then processed through DADA2 for ASV inference (Fig. S2). Default settings were used throughout the DADA2 pipeline, except the DADA inference functions `dadaF` and `dadaR` which used the `pool =‘pseudo’` argument, to increase the likelihood of identifying rare taxa. Consequently, the chimera removal function `removeBimeraDenovo` included the `method =‘pooled’` argument (Callahan *et al*., 2016b).

ASVs identified from the 16S rRNA gene amplicon data were assigned taxonomy using the Silva database (Yilmaz *et al*., 2013), and the quality of the data was assessed using the included controls (Fig. S5). Any ASVs identified as chloroplasts, mitochondria, or archaea, were subsequently removed from the data. Test Phase *Brassicaceae* 16S rRNA sequencing data was subsequently re-analysed independently following the described protocol to avoid any biases from the six no-template negative controls, and the four mock communities. These are the Test Phase *Brassicaceae* ASVs which are reported hereafter.

### α-diversity of the Test Phase *Brassicaceae* rhizosphere and root communities

First, to identify any changes in abundance of the bacterial ASVs within the Test Phase *Brassicaceae* species, we estimated the absolute abundance of the bacterial 16S rRNA gene in each Test Phase DNA sample by qPCR (Azarbad *et al*., 2018; Props *et al*., 2017; see Supplementary Methods for details, Fig. S6). This additional data allowed us to refine our analyses to better contextualise the dynamics of the bacterial communities, found among the rhizospheres and roots of the five *Brassicaceae* crop species.

Second, to visualise taxonomic diversity, ASVs were plotted as taxa cluster maps using `heat_tree` from the `metacoder` package (Foster *et al*., 2017) for the rhizosphere and roots of both sampling years, where nodes represent phyla and class: node colours represent the absolute abundance of each ASV, while node size indicates the number of unique taxa. Taxa cluster maps facilitate visualizing abundance, as well as diversity across taxonomic hierarchies (Foster *et al*., 2017). Relative and absolute abundance bar charts were plotted at the phyla level for comparison. Finally, in order to estimate the coverage of the bacterial domain of life, we incorporated phylogenies into the phyloseq object following the method described by Callahan *et al*., 2016b (see Supplementary Methods for details). Faith’s phylogenetic diversity was calculated as an α-diversity index from the Test Phase *Brassicaceae* samples using the `pd` function from the `picante` package (Kembel *et al*., 2010; sum of all branch lengths separating taxa in a community). For comparison, Simpson and Shannon’s α-diversity indices were calculated (Fig. S7). Log transformed phylogenetic diversity indices were confirmed to respect normality (see Supplementary Methods for details).

We assessed differences between the mean phylogenetic diversity between soil histories, *Brassicaceae* hosts, and their interactions, using a Multi-Factor ANOVA and Tukey’s Post-Hoc test for significant groups that respected the assumptions of normality (Azarbad *et al*., 2020, Wang *et al*., 2020; see Supplementary Methods for details). Where normality could not be respected, we used the non-parametric Kruskal-Wallis rank sum test, `kruskal.test`. Specific groups of statistical significance were identified with the post-hoc pairwise Wilcoxon Rank Sum Tests, `pairwise.wilcox.test`, with the FDR correction on the p-values to account for multiple comparisons. As the relative and absolute abundance datasets yielded similar α-diversities, with and without ASV rarefaction using `rarefy_even_depth` (McMurdie & Holmes, 2013), only the results incorporating the absolute abundance are reported.

### Identification of differentially abundant ASVs and specific indicator species

To refine our understanding of the abundance and composition of the Test Phase *Brassicaceae* bacterial communities, we used two complementary methods to identify taxa specific to soil histories, or *Brassicaceae* hosts (see Supplementary Methods for details). First, taxa cluster maps were used to calculate the differential abundance of ASVs between experimental groups, including rhizosphere and root compartments, *Brassicaceae* host plants, and soil histories. Second, indicator species analysis was used to detect ASVs that were preferentially abundant in pre-defined environmental groups (soil histories, or *Brassicaceae* host). A significant indicator value is obtained if an ASV has a large mean abundance within a group, compared to another group (specificity), and has a presence in most samples of that group (fidelity) (Legendre & Legendre, 2012). The fidelity component complements the differential abundance approach between taxa clusters, which only considers abundance. Moreover, given the large number of taxa in our study, it is not practical to view taxa clusters as matrices below class, whereas indicator species analysis pinpoints specific ASVs of interest.

### β-diversity of the Phase 2 *Brassicaceae* rhizosphere and root communities

To test for significant community differences between both experiments, compartments, soil histories and *Brassicaceae* hosts, we used the non-parametric permutational multivariate ANOVA (PERMANOVA), where any variation in the ordinated data distance matrix is divided among all the pairs of specified experimental factors. The PERMANOVA was calculated using the `adonis` function in the `vegan` package (Oksanen *et al*., 2020), with 9999 permutations, and the experimental blocks were included as “strata”. Our preliminary PERMANOVA (Table S3) used a distance matrix calculated with the Bray-Curtis formula, and tested the significance of the effects of soil history, *Brassicaceae* host, compartment and experiment (Fig. S9). This was complemented with a PERMANOVA for each experiment and compartment, that specifically tested treatments and hosts as experimental factors, and used a weighed UniFrac distance matrix (Lozupone & Knight, 2005; Lozupone *et al*., 2007). This distance index incorporates the phylogenetic relationship of each dataset and the absolute abundance of each ASV, as estimated by qPCR of the 16S rRNA gene (Lozupone & Knight, 2008).

We used a variance partition, as a complement to the PERMANOVA, to model the explanatory power of soil history, *Brassicaceae* host, and soil chemistry in the structure of the Test Phase *Brassicaceae* bacterial communities. We then quantified how each significant factor (ie, the explanatory variables) impacted bacterial community structure with a distance-based redundancy analysis (db-RDA) (Legendre & Legendre, 2012). First, singleton ASVs were removed before the p`hyloseq` data were transformed using Hellinger’s, such that ASVs with high abundances and few zeros are treated equivalently to those with low abundances and many zeros (Legendre & de Cáceres, 2013). A weighted UniFrac index was calculated (Lozupone & Knight, 2005; Lozupone *et al*., 2007), where this distance index gives an estimate of how similar communities contain more phylogenetically related ASVs, weighed by the absolute abundance of each ASV, as estimated by qPCR of the 16S rRNA gene (Lozupone & Knight, 2008). Using a weighted UniFrac index in an ordination provides one way to test if bacterial community composition follows the evolutionary history within the *Brassicaceae* host plant family, as determining community distances based only on the number of shared taxa does not account for evolutionary distances between taxa, which are often extremely diverse among microbes (Fitzpatrick *et al*., 2018; Walters *et al*., 2018). The weighted UniFrac distance matrix was calculated using the `distance` function in p`hyloseq` (McMurdie & Holmes, 2013), and gave similar results as a Bray-Curtis distance matrix.

With the `vegan` package (Oksanen *et al*., 2020), soil chemistry was standardized (Legendre & Legendre, 2012) using the `decostand` function. We modelled the explanatory power of each experimental factor in each compartment from both experiments with a variance partition of a partial RDA, using the `varpart` function, and the weighted UniFrac distance matrix (Borcard *et al*., 1992). Variation in the bacterial community data not described by the explanatory variables were quantified by the residuals. Finally, to quantify the amount of variation described by each explanatory factor, db-RDA were calculated using the `capscale` function, and plotted using p`hyloseq` (McMurdie & Holmes, 2013).

## Results

### The bacterial rhizosphere communities were larger and more diverse than the root communities

In order to estimate the composition of the bacterial rhizosphere and root communities from the Test Phase *Brassicaceae* species, we used the DADA2 pipeline (Callahan *et al*., 2016 & 2017) to infer the retained16S rRNA amplicons as ASVs. We retained 10 178 467 high-quality 16S rRNA MiSeq amplicons (43 129 ±18 032 reads/sample) through the pipeline (Table 1 & Fig. S4). More reads were retained in the Test Phase *Brassicaceae* root samples than from the rhizosphere in both experiments (Table 1). However, more bacterial ASVs were consistently identified in the rhizosphere compared to the root communities (Table 1). The absolute abundance, or size, of each Test Phase bacterial community from both experiments were estimated by qPCR amplification of the 16S rRNA gene, where we observed that the bacterial rhizosphere communities were consistently larger than the root communities (Table 1). Finally, the phylogenetic diversity was consistently larger in the Test Phase bacterial rhizosphere communities, compared to the root communities (Fig. 1); other α-diversity indices illustrated the same trend (Fig. S7).

**Table 1.** The bacterial rhizosphere communities had more unique ASVs and larger community sizes than the root communities of five *Brassicaceae* host plants in the Test Phase of a two-year crop rotation, harvested in 2016 and 2017 from Swift Current, Sask. Raw reads were produced via Illumina’s MiSeq at Génome Québec, and processed through DADA2, where 10 178 467 reads were retained (16S rRNA Reads reported here) for ASV inference. A total of 37 445 ASVs were identified across the entire dataset. Bacterial community size was estimated by qPCR as the number of copies of the 16S rRNA gene.

|  |  | 16S rRNA Reads | ASV Occurrence | 16S rRNA Gene Copies / Sample |
| --- | --- | --- | --- | --- |
| Experiment 1<br>Test Phase | Rhizosphere | 1 139 535<br>(18 992 ± 4333 / sample) | 14 047 | 25 747 358<br>± 17 838 649 |
|  | Root | 3 315 289<br>(55 254 ± 5600 / sample) | 3009 | 2 823 077<br>± 2 099 135 |
| Experiment 2<br>Test Phase | Rhizosphere | 2 122 186<br>(37 896 ± 7080 / sample) | 26 001 | 7 692 764<br>± 4 988 172 |
|  | Root | 3 601 457<br>(60 024 ± 11 835 / sample) | 4307 | 3 509 901<br>± 1 818 824 |

**Figure 1.**
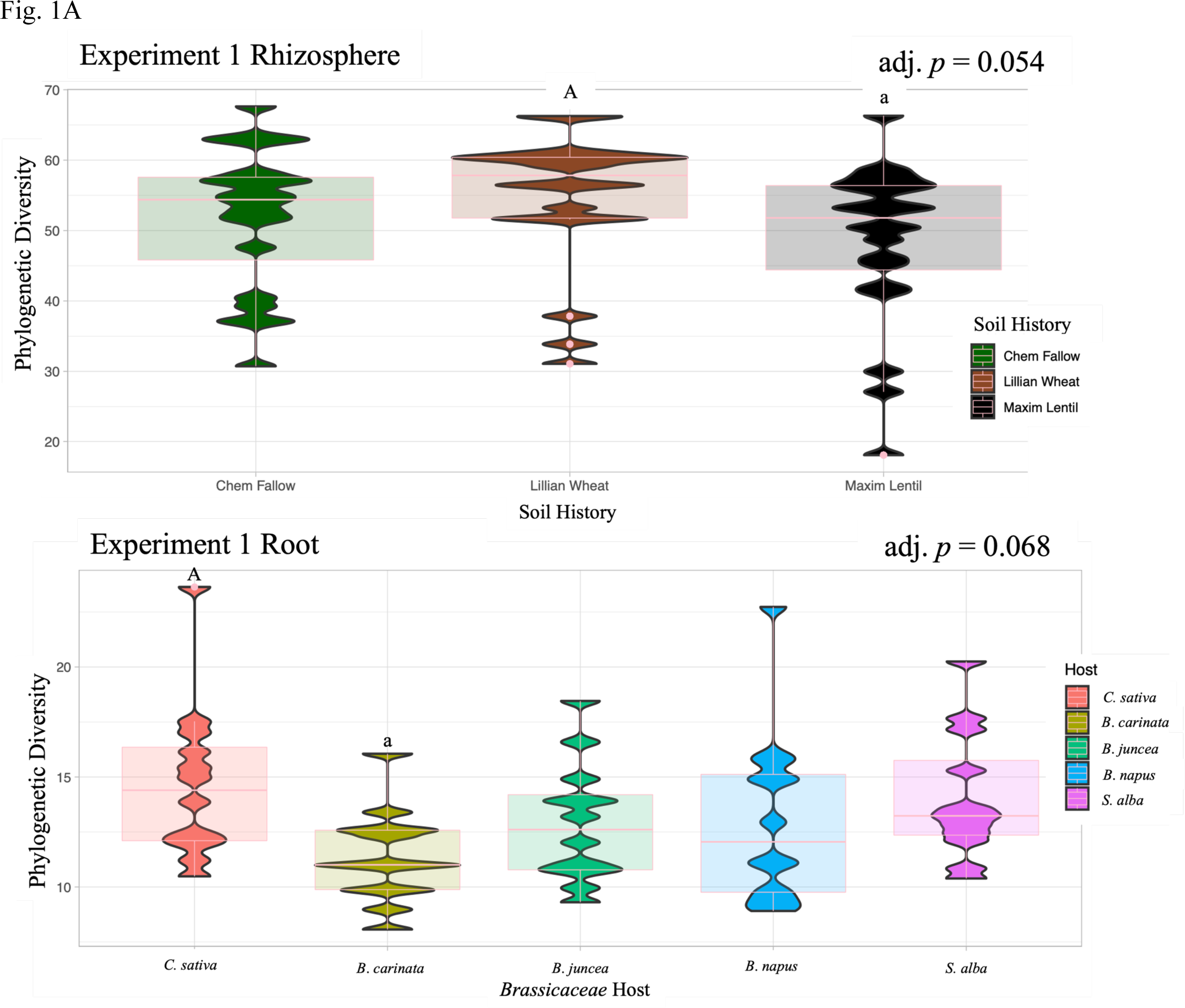
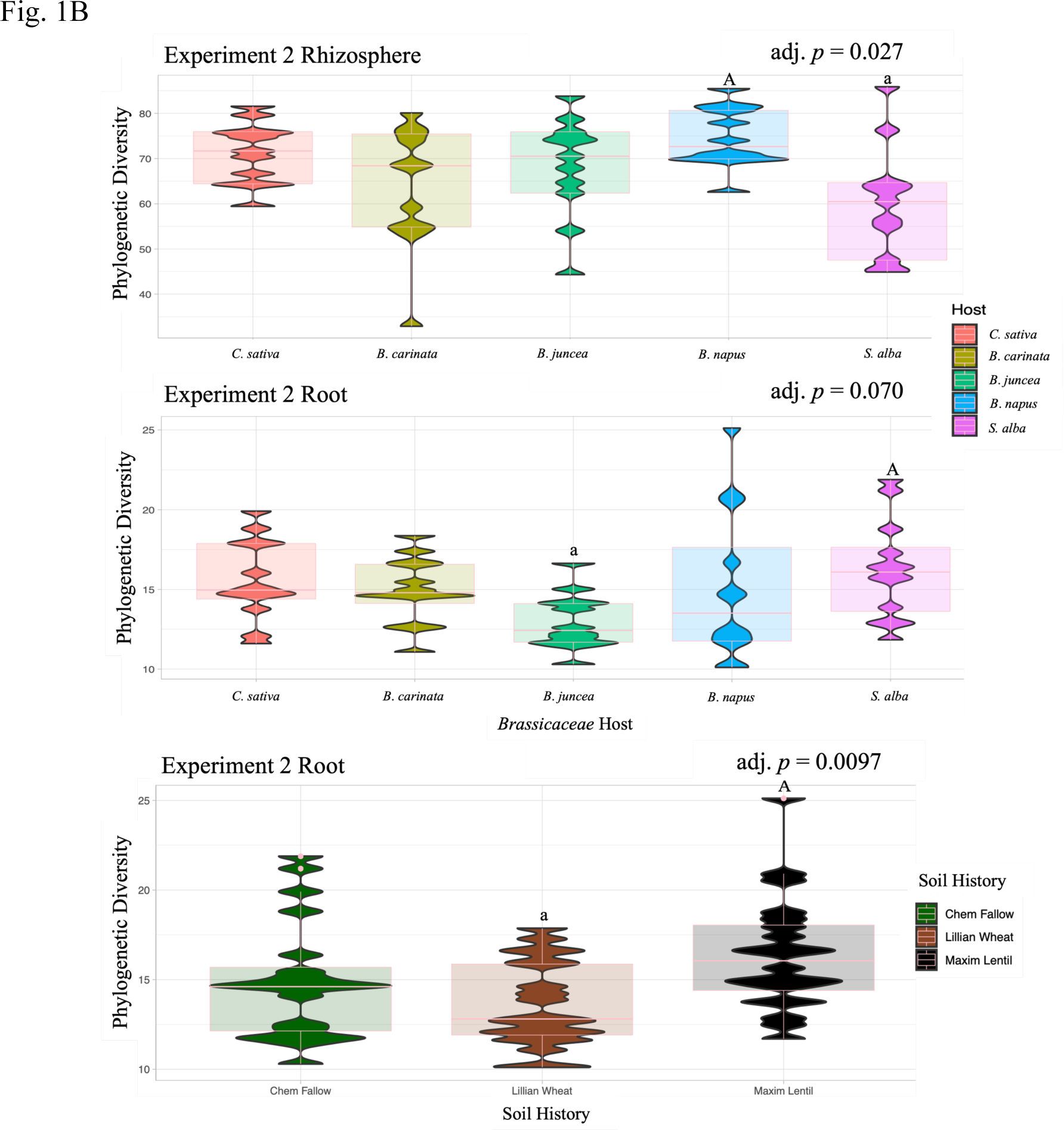
Phylogenetic diversity was consistently higher in the bacterial rhizosphere communities than in the root communities from five *Brassicaceae* host plants in the Test Phase of a two-year rotation, harvested in 2016 (A, Experiment 1) and 2017 (B, Experiment) from Swift Current, Sask. Phylogenetic diversity, the sum of branch lengths of a phylogenetic tree connecting all species in the sample, accounts for the wide evolutionary distance among bacterial populations, and better represents the functions present (Hsieh & Chao, 2017). Diversity indices were log-transformed, before being tested with a nested ANOVA, which confirmed the soil histories established in the first Conditioning Phase, and the Test Phase *Brassicaceae* hosts did not interact. Statistically significant groups were identified using Tukey’s post-hoc test. (A, top panel) Bacterial communities from the Test Phase rhizosphere with Conditioning Phase soil histories of growing wheat were significantly more (adj. *p* = 0.054) phylogenetically diverse than those communities with lentil soil histories. The non-parametric Kruskal test was used to test for significance among Test Phase communities grouped by Conditioning Phase soil histories. (A, bottom panel) Bacterial root communities from the Test Phase in Experiment 1 with *Camelina sativa* cv. Midas hosts were more phylogenetically diverse than root communities from *Brassica carinata* (adj. *p* = 0.0679). (B, top panel) Bacterial communities from the Test Phase rhizosphere from Experiment 2 with *Brassica napus* cv. canola hosts were significantly more (adj. *p* = 0.027) phylogenetically diverse than rhizosphere communities with *Sinapis alba* cv. Polish hosts. (B, middle panel) Experiment 2 bacterial root communities from Test Phase *Sinapis alba* cv. Polish hosts were more phylogenetically diverse than root communities from *Brassica juncea* cv. Cutlass hosts (adj. *p* = 0.070). (B, bottom panel) Bacterial communities from the Test Phase roots from Experiment 2 with Conditioning Phase soil histories of growing lentils were significantly more (adj. *p* = 0.0097) phylogenetically diverse than those communities with wheat soil histories.

ASVs were plotted as taxa clusters to the class level, where we observed that the bacterial communities from Experiment 1 (Fig. 2) and 2 (Fig. S8A) were dominated by phyla *Acidobacteria* (classes *Acidobacteria* and *Blastocatellia*), *Actinobacteria* (*Actinobacteria*), *Bacteroidetes* (Bacteroidia), *Firmicutes* (*Clostridia*), *Proteobacteria* (*Gammaproteobacteria*), and *Verrucomicrobia* (*Verrucomicrobae*). These taxa were dominant in both the rhizosphere and root communities (Fig. 2 & Fig. S8A). Moreover, we observed that all the taxa identified in the root communities, were also present in the rhizosphere.

**Figure 2.**
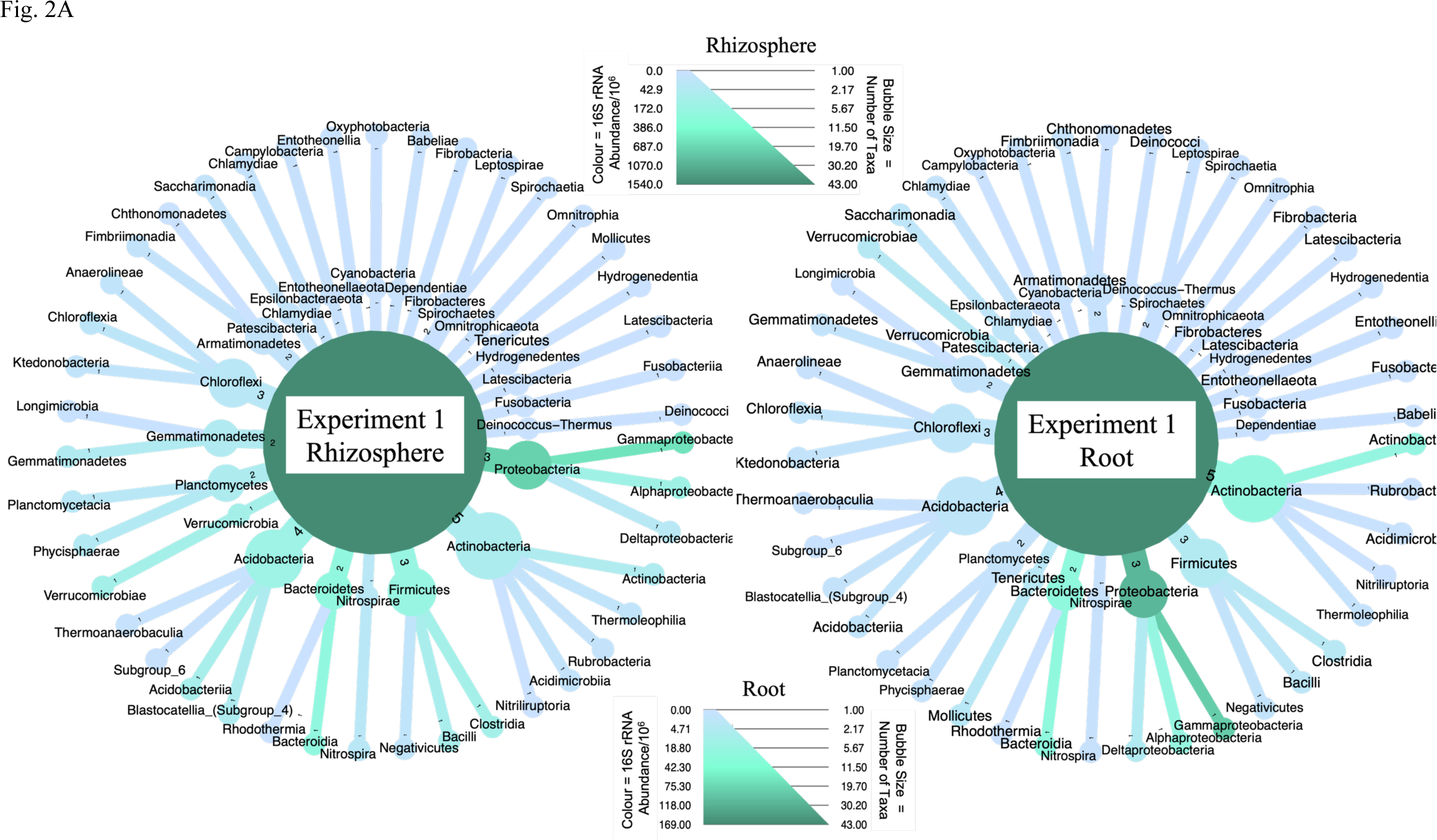
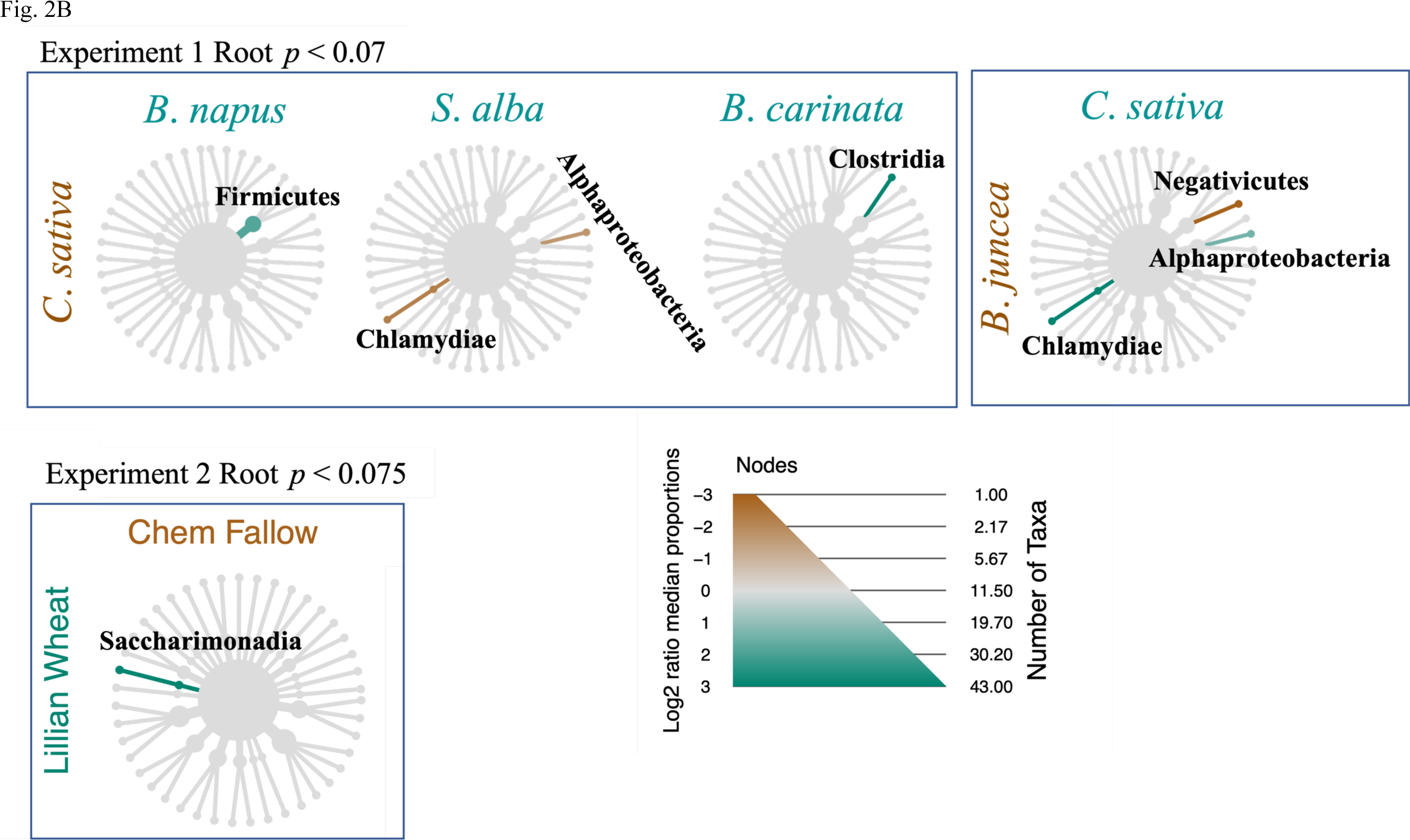
The abundance and composition of the bacterial communities varies noticeably between the rhizosphere (A, left) and root (A, right), as represented by taxa clusters of ASVs inferred from among the five *Brassicaceae* host plants in the Test Phase of a two-year rotation, harvested in 2016 from Swift Current, Sask. ASV absolute abundance, represented by the colour scale, was estimated by multiplying the 16S rRNA copy number / 10^6^ by the ASV abundance, while the size of each bubble represents the number of unique taxa represented to the class level. (A, left) Experiment 1 Test Phase bacterial rhizosphere communities were larger (1.5 ×10^9^ ASVs) than the root communities (1.69 ×10^8^ ASVs, right). (B) Test Phase *Brassicaceae* host plants only had significant variation in terms of composition among their root bacterial communities; among the root communities in Experiment 1 there were only differences between *Brassicaceae* hosts (B, top panel, *p*. adj < 0.07), while among the root communities in Experiment 2 there were significant differences between soil histories (B, bottom panel, *p*. adj < 0.075). Significantly enriched taxa, labelled in bold, were tested between each pair of host plants and soil history. Taxa that were significantly more abundant are highlighted brown or green, following the labels for each compared host. These differential taxa clusters identified significantly enriched (ie, abundant), using the non-parametric Kruskal test, followed by the post-hoc pairwise Wilcox test, with an FDR correction.

### Soil history was only significant in structuring bacterial communities in the driest year

Next, we evaluated our hypothesis that the three soil histories established the previous year would impact the structure of these bacterial communities differently. Soil history significantly structured the Test Phase bacterial communities only in Experiment 2, in both the rhizosphere (PERM *R*^*2*^ = 0.0722, *p* = 0.0387) and root (PERM *R*^*2*^ = 0.0873, *p* = 0.003, Table 2). Soil history was also identified as a significant factor in Experiment 2 by variance partition, both in the Test Phase rhizosphere communities (0.4%, *p* < 0.01, Fig. 3A), and root communities (4.3%, *p* < 0.01, Fig. 3B). RDA further confirmed soil history significantly structured the bacterial communities in Experiment 2. The Test Phase root communities followed a clear gradient between the three soil histories (adj. *R*^*2*^ = 0.0989, *p* = 0.001, Fig. 3C). The relationship among the Test Phase rhizosphere communities and soil history remained significant, though a similar gradient between the three soil histories was less clear (adj. *R*^*2*^ = 0.0399, *p* = 0.019, Fig. S11A).

**Table 2.** PERMANOVA identified that the *Brassicaceae* host plants were always a significant experimental factor in structuring the bacterial rhizosphere and root communities from the Test Phase of a two-year crop rotation, harvested in 2016 (Experiment 1) and 2017 (Experiment 2) from Swift Current, Sask. The previous year’s soil history was only significant in the Test Phase bacterial rhizosphere and root communities of Experiment 2, while the *Brassicaceae* host ∼ crop history interaction was never significant. The PERMANOVA was calculated using a weighted UniFrac distance matrix, with 9999 permutations.

| A | Experiment 1 |  |  |  |  |  |
| --- | --- | --- | --- | --- | --- | --- |
|  | Rhizosphere |  |  | Root |  |  |
|  | F Model | R<sup>2</sup> | Pr (> F) | F Model | R<sup>2</sup> | Pr (> F) |
| Soil History | 1.859 | 0.0524 | 0.0789 | 1.271 | 0.0332 | 0.2207 |
| *Brassicaceae* Host | 3.912 | 0.2207 | **0.0002** | 5.267 | 0.2751 | **0.0001** |
| Soil History *<br>*Brassicaceae* Host | 0.820 | 0.0925 | 0.7603 | 0.995 | 0.104 | 0.4577 |
| B | Experiment 2 |  |  |  |  |  |
|  | Rhizosphere |  |  | Root |  |  |
|  | F Model | R<sup>2</sup> | Pr (> F) | F Model | R<sup>2</sup> | Pr (> F) |
| Soil History | 2.1358 | 0.0722 | **0.0387** | 2.7623 | 0.0873 | **0.0003** |
| *Brassicaceae* Host | 2.0705 | 0.1400 | **0.0097** | 1.7879 | 0.1130 | **0.0045** |
| Crop History *<br>*Brassicaceae* Host | 0.6999 | 0.0947 | 0.8860 | 0.7014 | 0.0887 | 0.9666 |

**Figure 3.**
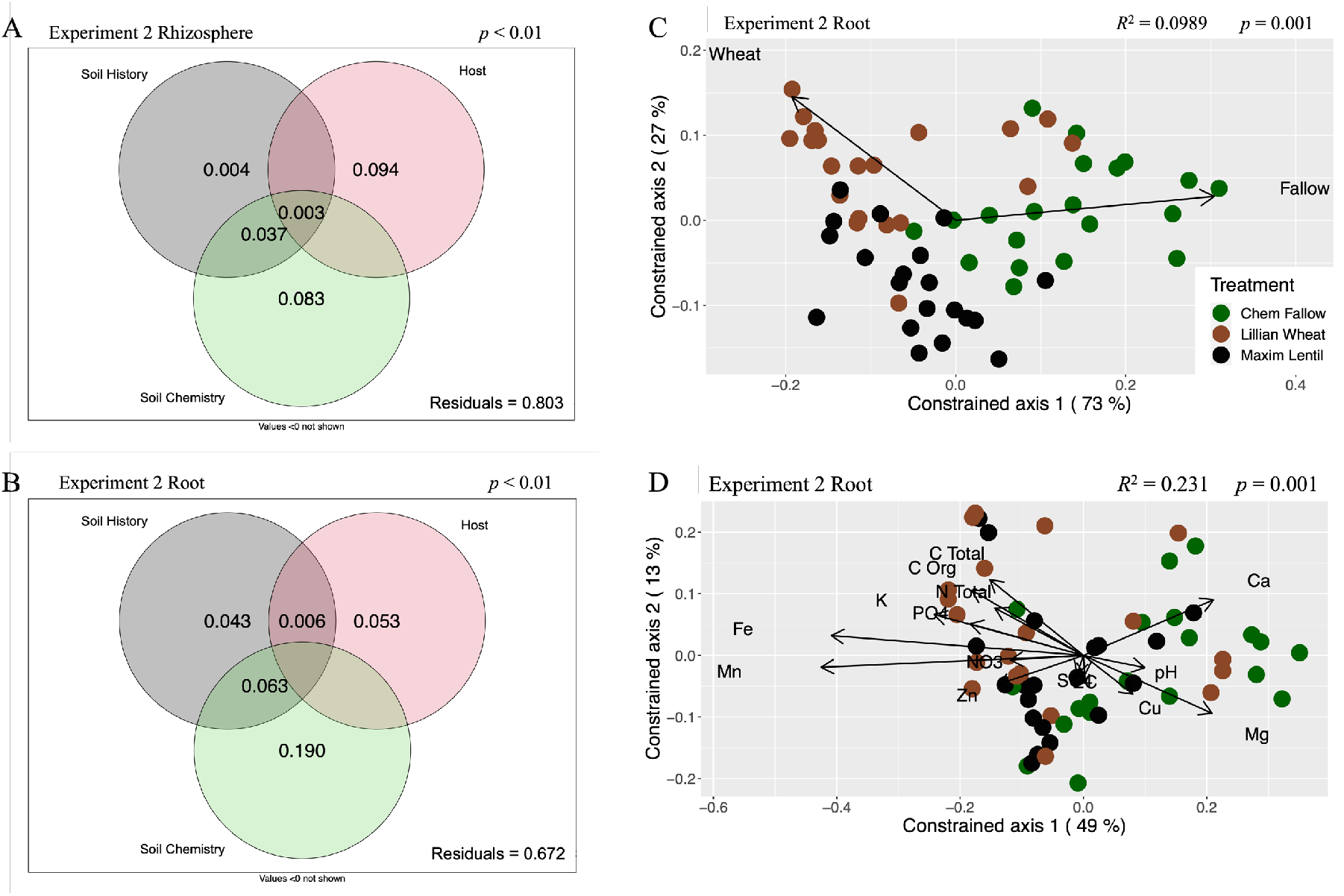
Soil chemistry explained the most variance in the bacterial community structures in the rhizosphere and roots of five *Brassicaceae* host plants in the Test Phase of a two-year rotation, harvested in 2017 from Swift Current, Sask. Weighted UniFrac distances were used with a variance partition (A & B), which modelled the explanatory power of each experimental factor (*Brassicaceae* host, soil history, and soil chemistry) in the Test Phase bacterial communities. Distance-based redundancy analyses (C & D) quantified how the experimental factors impacted community structure, where communities with similar phylogenetic composition appear closer together. (A) Variance partition illustrated the strong influence of *Brassicaceae* host plants (9.4%) and soil chemistry (8.3%) in explaining the Test Phase rhizosphere communities in Experiment 2. (B) The influence of soil chemistry increased (19%), as did soil history (4.3%) in the Test Phase bacterial root communities in Experiment 2. (C) Soil history was still significant (*R*^*2*^ = 0.0989, *p* = 0.001) in structuring the Test Phase bacterial root communities in Experiment 2, though soil chemistry was more explanatory (D, *R*^*2*^ = 0.231, *p* = 0.001). D) pH was opposed by potassium, as well as phosphate, while calcium was contrasted by zinc, and magnesium was contrasted by total nitrogen and organic carbon. However, soil chemistry does not have a clear relationship in explaining the phylogenetic similarity between communities, as a function of soil history, nor *Brassicaceae* host plant.

Soil history also had a significant impact on the composition of the Test Phase bacterial root communities in Experiment 2. First, root communities with a lentil soil history had significantly higher phylogenetic diversity (*p*. adj = 0.0097, Fig. 1B, bottom panel) than those root communities with wheat soil histories. Second, Test Phase root communities with wheat soil histories were enriched in *Patescibacteria* (*Saccharimonadia*), compared to those Test Phase root communities with fallow soil histories (*p*. adj < 0.075, Fig. 2B). Finally, indicator species analysis of Experiment 2 detected ASVs that were preferentially abundant in the Test Phase *Brassicaceae* roots with lentil or fallow soil histories (*p*. adj < 0.05, Table 3). Conversely, soil history did not significantly impact the size of the bacterial communities in Experiment 2, nor did soil history appear to impact the composition of the Test Phase rhizosphere communities in the experiment (Fig. S8A). Nonetheless, our data illustrated an important role for soil history in structuring the Test Phase bacterial communities in Experiment 2, especially the root communities.

**Table 3.** Indicator species were identified in the bacterial root communities in the Test Phase of a two-year crop rotation, harvested in 2016 (Experiment 1) and 2017 (Experiment 2) from Swift Current, Sask. In Experiment 1, seven ASVs were identified as indicator species in the Test Phase root communities of *Camelina sativa*. In Experiment 2, ten ASVs were identified from Test Phase root communities with soil history of growing lentils in the previous Conditioning Phase, while ten different ASVs were identified as indicator species from root communities with a fallow soil history. Indicator species analysis relies on abundance and site specificity to statistically test each ASV, which we report here as *p* < 0.05, with a FDR correction.

| Most closest taxon | Experiment 1<br>Roots ($p < 0.05$) | Experiment 2<br>Roots ($p < 0.05$) |
| --- | --- | --- |
| Alphaproteobacteria, *Phenylobacterium* (3x ASVs) | *C. sativa* |  |
| Actinobacteria, *Mycobacterium* | *C. sativa* |  |
| Gammaproteobacteria, Burkholderiales, *Aquabacteria* | *C. sativa* |  |
| Alphaproteobacteria, Micropepsaceae (2x ASVs) | *C. sativa* |  |
| Bacteroidetes, *Mucilaginibacter* |  | Lentil |
| Bacteroidetes, Chitinophagales, *Chitinophaga* (2x ASVs) |  | Lentil |
| Alphaproteobacteria, *Sphingomonas* |  | Lentil |
| Alphaproteobacteria, Rhizobales, *Labrys* |  | Lentil |
| Alphaproteobacteria, *Caulobacter* |  | Lentil |
| Gammaproteobacteria, Burkholderiales, *Rhizobacter* (2x ASVs) |  | Lentil |
| Actinobacteria, *Mycobacterium* |  | Lentil |
| Gammaproteobacteria, Xanthomonadales, *Luteibacter* |  | Lentil |
| Bacteroidetes, Chitinophagales, *Niastella* (4x ASVs) |  | Fallow |
| Gammaproteobacteria, *Pseudomonas* (3x ASVs) |  | Fallow |
| Gammaproteobacteria, Burkholderiales, *Massilia* |  | Fallow |
| Gammaproteobacteria, Xanthomonadales, *Lysobacter* |  | Fallow |
| Firmicutes, Bacilli, *Paenibacillus* |  | Fallow |

Soil history was comparatively weaker in structuring the Test Phase bacterial communities in Experiment 1 (rhizophere: PERM *R*^*2*^ = 0.0524, *p* = 0.0789; root: PERM *R*^*2*^ = 0.0332, *p* = 0.2207, Table 2). Variance partition, however, did identify soil history as significant (*p* < 0.001; Fig. 4A), though soil history only explained 1.6% of the Test Phase bacterial rhizosphere communities. RDA illustrated a gradient between Test Phase bacterial rhizosphere communities with fallow soil histories, versus communities that were conditioned with lentil, or wheat (adj. *R*^*2*^ = 0.027, *p* = 0.054, Fig. S10A). Phylogenetic diversity was also higher among Test Phase rhizosphere communities from Experiment 1 with wheat soil histories (*p*. adj = 0.054, Fig. 1A, top panel), compared to those rhizosphere communities conditioned with lentils. The only evidence we observed where soil history significantly influenced the Test Phase bacterial root communities in Experiment 1 was in the variance partition, where soil history explained 0.8% of the root communities (*p* < 0.001; Fig. 4B).

**Figure 4.**
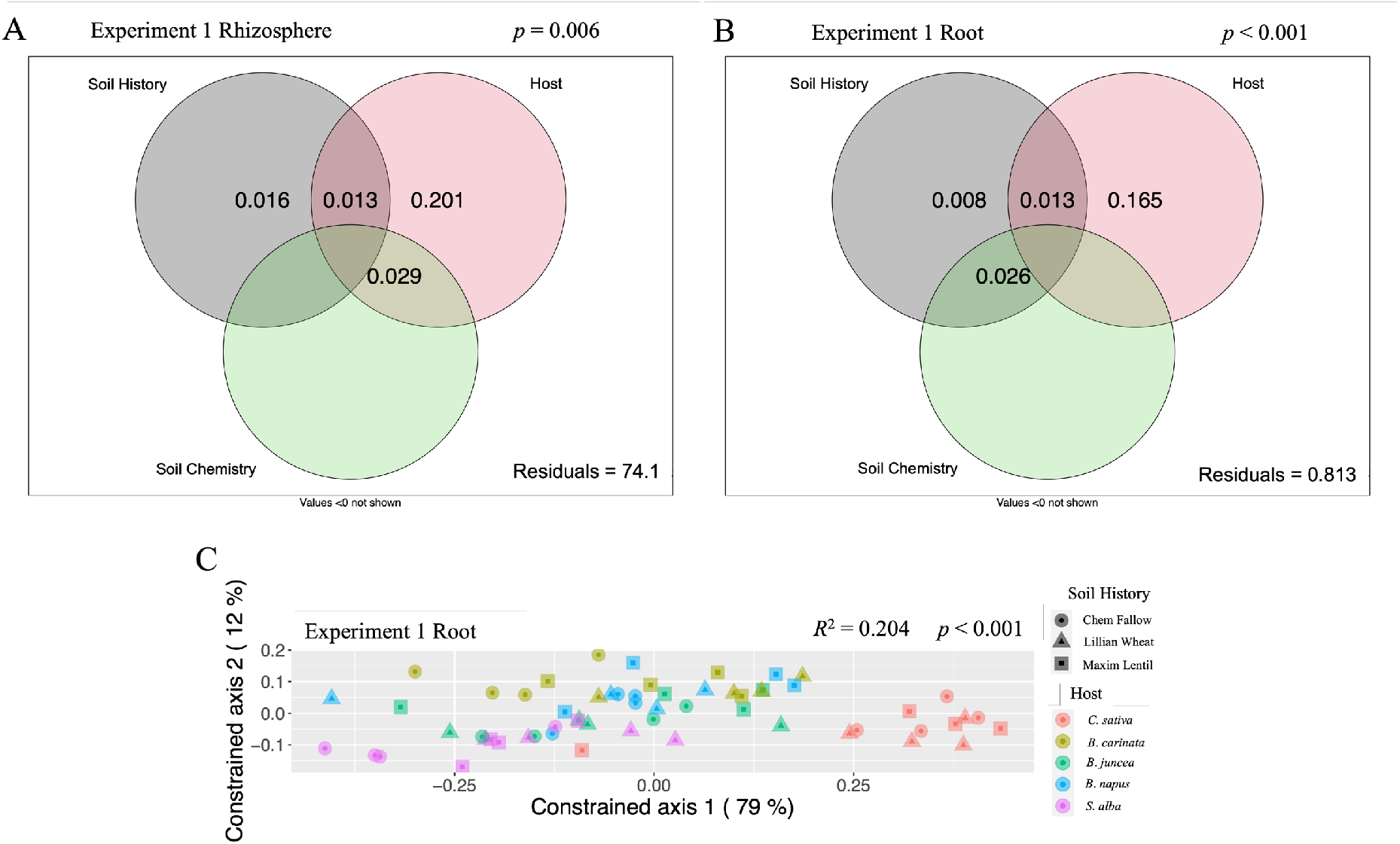
Bacterial communities were primarily structured by *Brassicaceae* host plants among rhizosphere and roots of five *Brassicaceae* host plants in the Test Phase of a two-year rotation, harvested in 2016 from Swift Current, Sask. Weighted UniFrac distances were used with a variance partition (A & B), which modelled the explanatory power of each experimental factor (*Brassicaceae* host, soil history, and soil chemistry) in the Test Phase bacterial communities of Experiment 1. Distance-based redundancy analyses (C) quantified how *Brassicaceae* host plants impacted community structure, where communities with similar phylogenetic composition appear closer together. Variance partition illustrated the strong influence of *Brassicaceae* host plants (20.1%) in explaining the Test Phase rhizosphere communities (A), and the root (B, 16.5%) communities in Experiment 1. (C) *Brassicaceae* host plants were also significant (*R*^*2*^ = 0.204, *p* < 0.001) in structuring the Test Phase bacterial root communities, as seen by RDA. *C. sativa* (red, at right) bacterial root communities were significantly more similar amongst themselves, than to root communities from other *Brassicaceae* host plants.

### Bacterial communities were influenced by soil chemistry in the driest year

The variance partition used to model the explanatory power of each factor in Experiment 2, revealed that the Test Phase bacterial communities were significantly explained by the soil chemistry (Fig. 3A & B). Soil history explained 0.4% of variance in the data among the rhizosphere communities, while soil chemistry accounted for 8.3% (Fig. 3A). RDA further supported the key role of soil chemistry in the rhizosphere communities of Experiment 2 (adj. *R*^*2*^ = 0.27, *p* = 0.001, Fig. S11B). Soil chemistry also explained the most variance in the Test Phase root communities of Experiment 2; 19% was attributed to soil chemistry, compared to 4.3% of the variance in the root community data being explained by soil history (Fig. 3B). RDA further supported the importance of soil chemistry in the root communities from the Test Phase of Experiment 2 (adj. *R*^*2*^ = 0.231, *p* = 0.001, Fig. 3D). These data strongly indicate that the Test Phase bacterial rhizosphere and root communities were strongly shaped by the soil chemistry in the dry year of Experiment 2, which was not detected in Experiment 1.

### *Brassicaceae* host plants were significant in structuring bacterial communities in both experiments

After identifying soil chemistry and soil history as key factors structuring the Test Phase bacterial communities in the dry year of Experiment 2, we evaluated our hypothesis that each of the five current *Brassicaceae* host plants would structure unique bacterial communities in their roots and rhizosphere. *Brassicaceae* host plants significantly structured the Test Phase bacterial communities, in both the rhizosphere and the root, in both Experiment 1 and 2 (Table 2). We noted there was never a significant interaction effect between soil history and *Brassicaceae* host plants, regardless of Experiment, or compartment (Table 2). The *Brassicaceae* host plants were also significant in the variance partition, and explained 9.4% and 5.3% of the community variance in the Test Phase rhizosphere and roots, respectively, of Experiment 2 (Fig. 3A & B). RDA illustrated a gradient between *C. sativa, B. carinata*, and *B. juncea* hosts in the Test Phase root communities in Experiment 2 (adj. *R*^*2*^ = 0.0307, *p* = 0.039, Fig. S12). RDA was not significant in the corresponding rhizosphere communities in Experiment 2, despite being significant in the root communities.

The *Brassicaceae* host plants also modified the composition of the Test Phase bacterial communities in Experiment 2. Rhizosphere communities from *B. napus* were more diverse than *S. alba* (*p*. adj = 0.0268, Fig. 1B). Experiment 2 root communities from *S. alba* hosts were significantly more diverse than *B. juncea* (*p*. adj = 0.0698, Fig. 1B). Furthermore, the mean absolute abundance of the root communities from *S. alba* were significantly smaller than root communities from *B. carinata* (pairwise Wilcox test, *p*. adj < 0.05, data not shown).

In Experiment 1, the *Brassicaceae* host plants were highly significant in structuring the Test Phase bacterial rhizosphere and root communities (Table 2). In the rhizosphere, the variance partition attributed 20.1% of the data as explained by the *Brassicaceae* hosts (Fig. 4A). Among the root community data, from Experiment 1, the variance partition modeled 16.5% of the data due to the *Brassicaceae* host plants (Fig. 4B). RDA illustrated how the bacterial rhizosphere communities from Experiment 1 were significantly structured by the *Brassicaceae* host (adj. *R*^*2*^ = 0.147, *p* = 0.0001, Fig. S10B). Rhizosphere communities from *C. sativa* were particularly impacted, as they appeared the most distinctly clustered. Interestingly, *B. carinata* had particularly diffuse communities (Fig. S10B), likely related to variation in species richness across the x-axis. The *Brassicaceae* host plants structured the Test Phase root communities from Experiment 1 even more drastically than their corresponding rhizosphere communities (adj. *R*^*2*^ = 0.204, *p* = 0.001, Fig. 4C); particularly for communities from *C. sativa*, which were distinctly clustered from communities with other host plants.

The *Brassicaceae* host plants created significant changes in the composition of the Test Phase root communities from Experiment 1, primarily in the *C. sativa* communities. First, we found that root communities from *C. sativa* were more phylogenetically diverse than those communities from *B. carinata* (*p*. adj = 0.0679, Fig. 1 bottom panel). *C. sativa* root communities were enriched in both *Alphaproteobateria* and *Chlamydiae*, compared to root communities from *S. alba* and *B. juncea* hosts, as determined by comparing taxa clusters of the bacterial root communities between each *Brassicaceae* host (*p*. adj < 0.07, Fig. 2B). *C. sativa* root communities were also enriched in *Firmicutes* (*Negativicutes*), compared to *B. juncea* hosts (Fig. 2B). However, *C. sativa* root communities were depleted in *Firmicutes* (*Clostridia*) compared to *B. carinata* communities, and were also depleted in *Firmicutes* compared to *B. napus* communities (Fig. 2B). Finally, indicator species analysis identified seven unique ASVs that were specific to the Test Phase bacterial root communities of *C. sativa* from Experiment 1 (Table 3). Taken together, our data points to a significant impact of *Brassicaceae* host plants on the structure of the Test Phase bacterial communities, especially in the roots of *C. sativa* from Experiment 1.

## Discussion

Previous soil history and the current plant host are plant-soil feedbacks (PSFs) that structure soil bacterial communities that operate over two time-scales (Hannula *et al*., 2021). We found that the current *Brassicaceae* host plants were the primary factor in structuring the bacterial rhizosphere and root communities in agricultural conditions, but that this influence could be weakened. We coupled 16S rRNA metabarcoding with a two-phase study design to test two hypotheses: 1) that each of the five current *Brassicaceae* host plants would structure unique bacterial communities in their roots and rhizosphere, and 2) that the three soil histories established the previous year would impact the structure of these bacterial communities differently.

We found that the current *Brassicaceae* host plants were always significant in structuring the Test Phase bacterial communities in two experiments done over two successive years that largely differed by the amount of rain received during the plant growth period. Moreover, the composition and diversity of the Test Phase bacterial root communities from the *C. sativa* host were particularly distinct compared to root communities from the other four *Brassicaceae* host plants, (2) the previous soil history established in the Conditioning Phase still had a significant effect on structuring the Test Phase bacterial rhizosphere and root communities a year later in Experiment 2, that was done on the dry year. In fact, our results illustrated that the Test Phase soil chemistry explained the most variance in the structure of the Test Phase bacterial communities in Experiment 2, while soil history and *Brassicaceae* host plants were secondary. In Experiment 1, the wet year, the previous soil history and the current soil chemistry had no effect on the Test Phase bacterial communities.

### Bacterial communities among *Brassicaceae* hosts

Our results from Experiment 1 illustrated that the influence of the PSF from the current *Brassicaceae* host plants was strong enough to obscure any influence from the soil history established from the previous Conditioning Phase (Fig. 4). In Experiment 2, the effect of *Brassicaceae* host plant, while still significant, was noticeably weaker (Fig. 3). Our results in both experiments illustrated that four of the *Brassicaceae* host plants, *S. alba, B. carinata, B. juncea* & *B. napus*, assembled essentially the same bacterial rhizosphere and root communities, over-riding any residual effects from soil history, and variation in the *Brassicaceae* management practices, and agricultural inputs. Conversely, *C. sativa* assembled significantly different bacterial communities, most prominently in the root communities.

Traditionally, there has been robust debate surrounding the degree to which host plants play a role in shaping their microbial communities (Erlandson *et al*., 2018). More recently, there are a growing number of studies that demonstrate stable bacterial relationships with host plants conserved through evolutionary time. Stopnisek & Shade (2021) reported that 48 bacterial taxa were persistent in *Phaseolus vulgaris* root communities across time and space, including between soils of different continents. Zhang *et al*., (2019) reported clear differences between *Oryza sativa* L. *indica* and *japonica* cultivars, and demonstrated that the SNP variation of a rice nitrate transport gene, *NRT*.*1B*, was responsible for enriching the root microbiome in bacterial nitrogen-use genes. Fitzpatrick *et al*., (2020) quantified larger host impacts on the bacterial root community than in the rhizosphere, for 30 angiosperms grown in a common soil. Finally, within the same bog, Wicaksono *et al*., (2021) identified similar bacterial communities, which were differentially enriched depending on if the host was a bryophyte, or a vascular plant. Thus, we see a variety of examples, through different plant taxonomic relationships, that demonstrate the important role of plant hosts in structuring their bacterial communities. Our results in comparing five *Brassicaceae* host plants in an agricultural setting fits with this perspective of host-bacteria interaction through evolution.

Detecting the influence of plant hosts on structuring their bacterial communities may require specific details worth highlighting. First, plants grown in soils from the same source, like a “Common Garden” design, will all experience the same environmental filter, and establishes a common microbial starting point (Bouffaud *et al*., 2014). Second, detecting the impact of the host plant will be easier when the hosts are at larger evolutionary distances. Wicaksono *et al*., (2021) demonstrated clear host dominance over soil between bryophytes and vascular plants. Similarly between angiosperm families, Fitzpatrick *et al*., (2020) had sufficient evolutionary distance between host plants for clear resolution. Detecting a clear impact of host plants within a given plant family, as we have reported here with our novel approach in the *Brassicaceae*, may create challenges; the evolutionary distance of the host plants may be insufficient to be detectable between the bacterial soil communities (Bouffaud *et al*., 2014). Future studies ought to design experiments as though designing a phylogenetic tree, and include a diversity of host plants, at varying evolutionary distances from the hosts of interest—as if adding appropriate outgroups (Bouffaud *et al*., 2014). In our study, *C. sativa* functioned as the outgroup to help identify any specific host plant impact on the bacterial communities from among the genus *Brassica*. Despite variation in genotypes, and agricultural management practices and inputs, we still observed *S. alba, B. carinata, B. juncea*, and *B. napus*, had phylogenetically similar bacterial communities. Conversely, we found phylogenetically distinct bacterial communities associated with *C. sativa*, particularly within the roots (Fig. 4). We observed a clear trend of *C. sativa* having compositional differences compared to the other *Brassicaceae* hosts (Figures 1, 2 & Table 3).

### Soil history and soil chemistry are revealed when the host plants feedback is weakened

Our results illustrated that the soil history established in the Conditioning Phase was still significant a year later in structuring the Test Phase bacterial communities in Experiment 2 (Fig. 3). Most striking in Experiment 2 is the division of phylogenetically distinct communities that are formed between root communities with different soil histories (Fig. 3C & Table 3). In Experiment 1, however, the PSF from the current *Brassicaceae* hosts dominated the community structure, and the influence of the PSF from the previous soil history was minimized (Fig. 4). From these results, we can observe how a novel PSF is established in the Conditioning Phase from the interaction of edaphic factors, the resident microbial community, and the presence, or absence, of specific resident plants, and their variation in agricultural treatments (Bouffaud *et al*., 2014). The three soil histories remained influential a year later in the Test Phase of Experiment 2, when the subsequent *Brassicaceae* host plants were planted and began to adjust the environmental screen. In Experiment 2 both scales of PSFs, the previously established soil history, and the current *Brassicaceae* host plants, were significant factors in structuring the bacterial communities (Table 2).

That both soil history and the *Brassicaceae* hosts were significant in Experiment 2, but not Experiment 1, is an interesting observation, as it suggests that the dominating, homogenizing effect that the *Brassicaceae* hosts had on structuring their bacterial communities was curtailed. In Experiment 1, the *Brassicaceae* host plants dominated the structure of the bacterial communities, to the exclusion of any effect from different soil histories, or any variation in soil chemistry due to the particular agricultural management practices, and inputs that were employed. Conversely, in Experiment 2, we observed the effect of the soil history that had been established the previous year in the Conditioning Phase, as well as a much-reduced effect from the *Brassicaceae* host plants. This could suggest that, despite identical experimental protocols, and the use of the appropriate agricultural management practices, and inputs, that another factor reduced the impact of the *Brassicaceae* host plants in Experiment 2.

One hypothesis for the disparate observations between Experiments 1 and 2, could be that the environmental conditions were 6x drier in the Test Phase of Experiment 2. If the *Brassicaceae* host plants were restricted in growth capacity (Fig. S3) due to the dry conditions, they would also be constrained in nutrient uptake from the soil, and rhizodeposition, as water availability is a key determinant of plant performance (Fitzpatrick *et al*., 2018). Our Experiment 2 data illustrated that soil chemistry accounted for the largest portion of the variance among the bacterial rhizosphere and root communities (Fig. 3). In our agricultural setting, soil chemistry is largely composed of the soil history, the agricultural management practices of the current and previous years, and the uptake and turnover from the *Brassicaceae* host plants, and microbial communities.. If the host plants were restricted in their capacities, it could account for their weakened impact on the bacterial communities, which was not observed in Experiment 1.

Drought conditions have been previously observed to alter bacterial soil communities. For example, Santos-Medellín *et al*., (2021) noted that various taxa of *Actinobacteria* were enriched in rice endosphere communities, as had been observed previously (Naylor *et al*., 2017, Santos-Medellín *et al*., 2017). Fitzpatrick *et al*., (2018) went a step further and noted that across multiple plant families, the enrichment of *Actinobacteria* appears adaptive for drought conditions. Our results in Experiment 2 also exhibited an enrichment in the absolute abundance of *Actinobacteria* (Fig. 2 vs Fig. S8), further suggesting that the *Brassicaceae* host plants were experiencing a drought event.

Previous work by Hannula *et al*., (2021) demonstrated that the impact of soil history on bacterial communities should fade rapidly. This is similar to our observations in Experiment 1, where the influence of the previous soil history had been entirely masked by the current *Brassicaceae* host plants. Conversely, if the *Brassicaceae* host plants in Experiment 2 were being limited due to adverse conditions, it may explain why soil history was still able to influence the bacterial soil communities. Our results, along with others (Schlaeppi *et al*., 2014; Dombrowski *et al*., 2016; Erlandson *et al*., 2018; Vieria *et al*., 2020), demonstrate the influence and interactions between soil history, host plants, and the current soil chemistry, in structuring the soil bacterial community.

### Benefits

In this study, we took advantage of an agricultural setting to test two hypotheses relevant to soil bacterial ecology, the time-scales involved in PSFs, and how previous soil history, and the current host plants, impact soil bacterial communities. Agricultural systems bridge the gap between controlled greenhouse conditions, and experiments in “natural” environments, and provide the opportunity to develop applications from our findings. For example, our data demonstrated that the PSF from the previous year’s soil history had a very limited impact, if any, on the soil bacterial communities the following year. This should limit concerns that these rotations may negatively disrupt the soil bacterial communities, at least in the short term, and encourage their use to help protect against potential fungal pathogens. Furthermore, our observations that four *Brassicaceae* hosts had largely similar bacterial communities, despite different previous soil histories, or current hosts, will be informative for further agricultural development, including breeding, rotations, or intercropping. Finally, that only the *C. sativa* host plants formed phylogenetically distinctive soil bacterial communities, is a key finding. It will be important to determine how and why this occurs, as well as if this distinct bacterial community is responsible for lower yields, and higher potential for the loss of nitrogen through denitrification, as reported by Wang *et al*., (2020).

### Conclusion

We found that the current *Brassicaceae* host plants, and the soil history established the previous year, both had PSF roles in structuring the bacterial communities of the roots and rhizosphere. The *Brassicaceae* host plants were consistently significant in structuring the bacterial communities, suggesting that the current plants quickly override any residual PSFs from previous soil history. Four host plants (*Sinapis alba, Brassica napus, B. juncea, B. carinata*) formed nearly the same bacterial communities, regardless of soil history. However, *Camelina sativa* host plants structured phylogenetically distinct bacterial communities compared to the other hosts, particularly in their roots. These are novel findings and illustrate that the development of these oilseed species has had limited impact on altering their bacterial communities. Furthermore, soil history established the previous year was only significant for bacterial community structure when the influence of the *Brassicaceae* host plants was weakened. From the evidence, we argued that some host plants experienced drought conditions, which negatively impacted their feedback on the soil, and suggest that this be further studied. Understanding how PSFs at different time-scales are involved in structuring bacterial communities is a limitation for employing microbiome technologies in improving agricultural systems. Our study highlights how PSFs and agricultural systems impact soil bacterial biodiversity, and offers new pathways forward for future research.

## Acknowledgements

This work was supported by the Natural Sciences and Engineering Research Council of Canada, CRD Fund (Grant Number: CRDPJ 500507-16), Canola Council of Canada and Saskatchewan Pulse Growers, which are gratefully acknowledged. We thank Yantai Gan for setting up and managing field experiments at Swift Current. We also thank Stéphane Daigle for assistance in statistical analysis, Chih-Ying Lay, Jacynthe Masse, Chantal Hamel, and Simon Joly and for their helpful comments and discussions. Finally, AB would like to thank Simon Morvan, Alexis Carteron, and the Quebec Centre for Biodiversity Science for teaching me to use R, and all their support in this project.

## Author Contributions

AJCB prepared the libraries for sequencing, performed the qPCR experiment, analyzed the data, and wrote the manuscript with input from all co-authors. LB conducted field Experiments 1 and 2 and collected data; MSA, & MH designed the experiment, supervised the work, contributed reagents, analytical tools, and revised the manuscript.

### Data Accessibility

Sequencing data and metadata are available at NCBI Bioproject under accession number: PRJNA786355.

