## Supplemental Materials for "What’s past is past, mostly: *Brassicaceae* host plants mask the feedback from the previous year’s soil history on bacterial communities, except when the *Brassicaceae* hosts experience drought"

### Supplementary Materials

#### Supplementary Methods

##### 16S rRNA gene amplicon generation

The 16S PCR reactions consisted of 11.5 μL $dH_2O$, 5.0 μL of 10X Buffer (Qiagen, Canada), 2.5 μL of 10 μM S-D-Bact-0341-b-S-17 forward and S-D-Bact-0785-a-A-21 reverse primers, commonly referred to as 341F and 805R, respectively (Alpha DNA, Montréal, Canada; Klindworth *et al*., 2012), 1.0 μL of dNTPs (Qiagen, Canada), and 0.5 μL of *T. aq* polymerase (Qiagen, Canada), for a total volume of 25 μL. 16S rRNA gene primers, F17 and R21, indicated in Supplementary Table 2, were synthesized with the CS1 and CS2 adapters, respectively, and HPLC purified, as per Genome Québec's (Montréal, Canada) submission requirements. PCR amplification of a 416 bp fragment from the V3-V4 region of the 16S rRNA gene (Klindworth *et al*., 2012) was run in an Eppendorf Mastercycle ProS (Germany) thermocycler, and consisted of an initial denaturation of 2 minutes at 95°C, followed by 30 cycles of 30 seconds denaturation at 95°C, 30 seconds annealing at 55°C, and 1 minute elongation at 72°C, before a final elongation of 5 minutes at 72°C (Bell *et al*., 2016; Lay *et al*., 2018).

##### Quality control of the 16S rRNA gene amplicon MiSeq data

In order to evaluate potential biases, or flaws, in our lab preparation, sequencing, and bioinformatic analysis, we included six no-template negative controls in our pipeline, two during DNA extraction, four during 16S rRNA gene amplification, as well as four replicates of a bacterial mock community of known composition (Fig. S2). All six no-template negative controls did yield reads, to a maximum of 3750 in the 2017 rhizosphere extraction sample (Fig. S4A), though they were all confirmed to contain no detectable DNA at any step during our lab manipulation. Given the drastic difference in the number of reads between the six no-template negative controls (Fig. S4A), and the replicates of the mock communities (Fig. S4B), we would suggest that the inferior numbers are sequencing artefacts. Moreover, the stringency of the `filterAndTrim` step eliminated nearly all the reads from these six samples (Fig. S4A).

The replicates of the bacterial mock community (Fig. S4B) also help to assess the accuracy and sensitivity of our lab preparation, sequencing, and particularly the bioinformatic analysis. The four samples were sequenced to an average of ~101 000 reads (Fig. S4B), suggesting a consistency in sequencing. Furthermore, Fig. S4B illustrates the reproducibility of the DADA2 pipeline, as each replicate looks similar through each step of the pipeline. Approximately a third of the reads were retained overall, which follows recommended standards (Callahan *et al*., 2016). The greatest loss of reads is consistently among the filtering step, further illustrating the stringency of this step to retain only high-quality reads for subsequent analysis.

Evaluating the inference of DADA2, the six no-template negative controls contained between 1 and 32 different ASVs identified from the few remaining reads. The majority were inferred as a single ASV identified to the phylum *Cyanobacteria* (Order Chloroplast), which we insured to have removed from the experimental Year 2 *Brassicaceae* data presented (Fig. S5A). The four mock community replicates again largely resembled each other in ASV composition. DADA2 inferred 34– 55 individual ASVs from the retained MiSeq reads, from a community composed of 20 possible ASVs (Fig. S5B). Highlighting the accuracy of the DADA2 pipeline to correctly infer ASVs, every bacterial species included in the mock community (Table S1) was detected in all four replicates.

The *Bacteroidetes*, *Deinococcus*, and *Actinobacteria*, were the most specific, as only 1, or 2, species were included in the mock community, and were accurately detected in the pipeline in each replicate (Fig. S5B). Interestingly, both mock replicates from among the endosphere samples added a third ASV identified as an *Actinobacteria*, while the other two replicates did not, despite there only being two *Actinobacteria* ASVs in the community. There was also an expansion among the ASVs identified as *Bacteriodetes* across all four replicates. Though there was only one *Bacteriodetes* included in the mock community, between five and ten were identified in the four mock replicates; the 2016 and 2017 endospheres contained six, the 2016 rhizosphere had 8, and the 2017 rhizosphere had 10. The mock community contained six *Proteobacteria*, yet eight, nine, 11, and 14, were identified in the 2016 rhizosphere, 2016 endosphere, 2017 endosphere, and 2017 rhizosphere, respectively. Fully half of the mock community was composed of *Firmicutes*, and from the possible ten included in the mock community, the 2016 rhizosphere and endosphere both identified 15 Firmicutes ASVs, while the 2017 rhizosphere and endosphere had 22, and 28, respectively (Fig. S5B). All of these "over-identifications" may be due to the heterogenous nature of colony inoculation. Finally, while there were no bacterial specimens of *Cyanobacteria*, nor *Chloroflexi*, included in the mock community, three of the mock replicates had an ASV identified as a *Cyanobacteria*, while one of the replicates also had an ASV identified as *Chloroflexi* (Fig. S5).

Two potential confounding issues with the 16S rRNA gene sequencing data from the Year 2 Brassicaceae bacterial communities are the depth of sequencing, and batch effects from sequencing. First, to confirm that the depth of Illumina sequencing was appropriate to detect the majority of bacteria present in each sample, the data was rarefied to the lowest number of retained reads, 2703, from among the all the Year 2 *Brassicaceae* samples and a rarefaction curve was plotted using the `vegan` package in R (Oksanen *et al.*, 2020; R Core Team, 2020). Most of the Year 2 *Brassicaceae* samples plateau by ~15 000 reads, and all by 20 000 reads, which suggests that all ASVs were estimated from the retained reads (43 129 ±18 032) in each samples.

To confirm that the Illumina MiSeq process did not introduce detectable biases into the data, an ordination was used to visual the distribution of the all the Year 2 *Brassicaceae* sequencing data (Fig. S9). Here we see that data falls into the most prominent groupings, compartment, rhizosphere, or root (Fig. S9A), and sampling year (Fig. S9B), as expected. Sequencing data produced from the same Illumina sequencing reaction that was biased in some way would be expected to group together in an unknown *a priori* fashion.

<u>Ordination</u>

As a preliminary exploration of the Year 2 data, the relative, and absolute abundance, were ordinated using a non-metric multidimensional scaling (NMDS) approach. First, singletons were removed from the data, which were subsequently transformed using the Hellinger transformation. The transformed data were then ordinated in `phyloseq` (McMurdie & Holmes, 2013), where the method was assigned as "NMDS", and the distance matrix was set as "bray". Similar results were obtained using principal co-ordinate analysis, and a Jaccard distance matrix. Ordinations were then plotted in `phyloseq`, and statistically tested using PERMANOVA, as described in the Methods section, except Year (2016 and 2017), and Compartment (Rhizosphere and Endosphere), were included in the model, and we used a Bray-Curtis distance matrix.

<u>Generating phylogenetic trees</u>

In order to employ phylogeny-based analysis methods, such as phylogenetic diversity, or UniFrac, for analyzing diversity of the Test Phase *Brassicaceae* bacterial communities, we assembled phylogenies for each compartment, from both sampling years, 2016 and 2017. Following the method described by Callahan *et al*., 2016b, 16S rRNA gene sequences for each ASV inferred from the Test Phase *Brassicaceae* data were aligned using a profile-to-profile algorithm (Wang & Dunbrack, 2004) with a dendrogram guide tree using the `decipher` package (Wright, 2016). With the `phangorn` package (Schliep, 2011), the maximum likelihood of each site was calculated using the `dist.ml` function using a JC69 equal base frequency model, before assembling phylogenies using the neighbour-joining method. An optimized GTR nucleotide substitution model was fitted to the phylogeny using the `optim.pml` function. Phylogenies were subsequently added to each `phyloseq` object.

<u>Estimating absolute abundance of bacterial communities by quantitative PCR</u>

In order to better understand and contextualise the dynamics of the bacterial communities found among the rhizospheres and roots of the five *Brassicaceae* crop species, we estimated the absolute abundance of the bacterial 16S rRNA gene in each Test Phase DNA sample by qPCR (Azarbad *et al*., 2018; Props *et al*., 2017). First, a standard curve of 16S rRNA gene copy numbers was constructed. Near full-length 1.5 kb 16S rRNA gene fragments were PCR amplified using the primers PA-27F-YM and PH-R (Bruce *et al*., 1992; Table S1) from DNA extracted from previously used soil samples (Lay *et al*., 2018). The PCR reaction and cycling conditions were as described above. The amplified 1.5 kb 16S rRNA gene fragment was visualized by a 0.7% gel electrophoresis, as described above, quantified using the QuBit dsDNA High Sensitivity Kit (Invitrogen, USA), and then serially diluted to $10^{-9}$. One µL of each dilution was then used as template in a 10 µL qPCR reaction.

The 16S rRNA gene qPCR reactions consisted of 5.0 µL of Maxima SYBR Green/ROX qPCR Mix (ThermoFisher Scientific, Canada), 3.4 µL $dH_2O$, 0.3 µL of 10 µM Eub 338 forward and Eub 518 reverse primers (Alpha DNA, Montréal, Canada; Fierer *et al*., 2005). All qPCR reactions were set-up in triplicate in 96-well plates using the Freedom EVO100 robot (Tecan, Switzerland), with a no-template negative control included on each plate. Reactions were run in a ViiA 7 Real-Time PCR System (Life Technologies, Canada) following the same cycling conditions as described previously for the 16S rRNA PCR amplification. The Eub338/Eub518 qPCR reaction amplified a 200 bp region of the V3 region (Bathe & Hausner, 2006; Davis *et al*., 2009; Muyzer *et al*., 1993). The number of 16S rRNA gene copies present in the serially diluted standard were calculated using the formula (Godornes *et al*., 2007):

$$\text{Number of 16S rRNA gene copies } \mu L^{-1} = \frac{\text{Avogadro's Constant x DNA } (g\ \mu L^{-1})}{\text{Number of base pairs x 600 Daltons}}$$

The standard curve for each diluted sample was plotted, with an $R^2$ value of 0.9938 and an amplification efficiency of -3.2013 (Fig. S6), falling within acceptable values (Fierer *et al*., 2005).

16S rRNA gene copy numbers were estimated for each Phase 2 sample by using 1 µL of a 1:10 dilution of DNA as template in the same 16S rRNA qPCR reaction and cycling conditions as described above for the standard curve. Melt curves generated by 0.5°C increments at the end of the qPCR programme confirmed amplicon specificity, and the 16S rRNA gene copy number was determined from the standard curve. A correction to determine the absolute abundance of ASVs from the Test Phase samples was achieved by multiplying the 16S rRNA gene copy number per ng, as estimated from the qPCR reaction, by the relative abundance matrix of ASVs identified (Azarbad *et al*., 2018; Bakker, 2018).

Total community absolute abundance per sample, and their distributions within each *Brassicaceae* host and Conditioning Phase soil history, were plotted. Absolute abundance data from the rhizosphere and root of each sampling year were log transformed. We used a Multi-Factor ANOVA to test for statistical differences in the means of community size between soil histories, *Brassicaceae* hosts, and their interaction, using the `anova` function (see Wang *et al.*, 2020 for details). All assumptions were confirmed to be respected: normality of the residuals established with a Shapiro-Wilk test, `shapiro.test`, while the heteroscedascity of residuals was confirmed with using a Bartlett test, `bartlett.test`. For significant ANOVAs, a post-hoc Tukey's Honest Significant Difference test, `TukeyHSD`, was used to determine which groups were statistically different.

<u>Identification of differentially abundant ASVs and specific indicator species</u>

Taxa cluster maps were generated using `compare_groups`, in the metacoder package (Foster *et al.*, 2017), the non-parametric Wilcoxon Rank Sum Tests determined if a randomly selected abundance from one group was greater on average than a randomly selected abundance from another group. As the statistical test was performed for each taxon, we used a false discovery rate (FDR) correction on the p-values to account for the multiple comparisons. When the comparison was between more than two groups, the differential abundances were plotted onto the taxa cluster map using `heat_tree_matrix` (Foster *et al.*, 2017).

We performed an indicator species analysis for the Test Phase ASVs identified in the rhizosphere and roots of both experiments. We obtained similar results from the relative, and absolute, abundance datasets, and so are only reporting the results using the absolute abundance. From the `indicspecies` package (De Cáceres & Legendre, 2009), we used the `multipatt` function with 9999 permutations. As the statistical test is performed for each ASV, we used the FDR correction on the p-values to account for multiple comparisons.

**Supplementary Tables**

**Table S1**. Bacterial strains included in the mock community (BEI Resources, USA) of known composition, was included on each plate (Fig. S2). The mock community contains DNA of 20 bacterial species in equimolar counts ($10^6$ copies/µL) of 16S rRNA genes. Taxa have been provided to illustrate the level of comparison.

| Bacteria | Taxonomy | | |
|---|---|---|---|
| | Phyla | Class | Order/Family |
| *Actinomyces odontolyticus* | Actinobacteria (P) | Actinomycetales (C) | |
| *Propionibacterium acnes* | Actinobacteria (P) | Actinomycetales (C) | |
| *Bacteroides vulgatus* | Bacteroidetes (P) | | |
| *Deinococcus radiodurans* | Deinococcus (P) | | |
| *Bacillus cereus* | Firmicutes (P) | Bacilli (C) | Bacillales (O)/ Bacillaceae (F) |
| *Listeria monocytogenes* | Firmicutes (P) | Bacilli (C) | Bacillales (O)/ Listeriaceae (F) |
| *Staphylococcus aureus* | Firmicutes (P) | Bacilli (C) | Staphylococcaceae (F) |
| *Staphylococcus epidermidis* | Firmicutes (P) | Bacilli (C) | Staphylococcaceae (F) |
| *Enterococcus faecalis* | Firmicutes (P) | Bacilli (C) | Lactobacillales (O)/ Enterococcaceae (F) |
| *Lactobacillus gasseri* | Firmicutes (P) | Bacilli (C) | Lactobacilliaceae (F) |
| *Streptococcus pneumoniae* | Firmicutes (P) | Bacilli (C) | Streptococcaceae (F) |
| *Streptococcus agalactiae* | Firmicutes (P) | Bacilli (C) | Streptococcaceae (F) |
| *Streptococcus mutans* | Firmicutes (P) | Bacilli (C) | Streptococcaceae (F) |
| *Clostridium beijerinckii* | Firmicutes (P) | Clostridia (C) | Clostridiale (O) |
| *Rhodobacter sphaeroides* | Proteobacteria (P) | Alphaproteobacteria (C) | |
| *Neisseria meningitides* | Proteobacteria (P) | Betaproteobacteria (C) | |
| *Helicobacter pylori* | Proteobacteria (P) | Epsilonproteobacteria (C) | |
| *Escherichia coli* K12 | Proteobacteria (P) | Gammaproteobacteria (C) | Enterobacteriales (O) |
| *Acinetobacter baumannii* | Proteobacteria (P) | Gammaproteobacteria (C) | Pseudomonadales (O)/ Moraxellaceae (F) |
| *Pseudomonas aeruginosa* PAO1-LAC | Proteobacteria (P) | Gammaproteobacteria (C) | Pseudomonadaceae (F) |

**Table S2**. Primers used in this study.

| Name | Sequence (5'-3') | Reference |
|---|---|---|
| S-D-Bact-0341-b-S-17 | CCTACGGGNGGCWGCAG | Klindworth *et al*., 2012 |
| S-D-Bact-0785-a-A-21 | GACTACHVGGGTATCTAATCC | Klindworth *et al*., 2012 |
| CS1 Adapters | ACACTGACGACATGGTTCTACA | Illumina, 2013 |
| CS2 Adapters | TACGGTAGCAGAGACTTGGTCT | Illumina, 2013 |
| 16S PA-27F-YM | AGAGTTTGATCCTGGCTCAG | Bruce *et al*., 1992 |
| 16S PH-R | AAGGAGGTGATCCAGCCGCA | Bruce *et al*., 1992 |
| Eub338 | ACTCCTACGGGAGGCAGCAG | Fierer *et al*., 2005 |
| Eub518 | ATTACCGCGGCTGCTGG | Fierer *et al*., 2005 |

**Table S3**. PERMANOVA for all the sampled Test Phase communities identified compartment (rhizosphere, or root), year harvested (2016 for Experiment 1, or 2017 for Experiment 2), *Brassicaceae* host, and soil history established in the Conditioning Phase, as significant experimental factors. PERMANOVA was calculated using a Bray-Curtis distance matrix, with 9999 permutations.

| A | Whole Dataset with Bray-Curtis Distances | | |
|---|---|---|---|
| | F Model | $R^2$ | Pr (> F) |
| Year | 25.170 | 0.07067 | **0.001** |
| Compartment | 68.750 | 0.19304 | **0.001** |
| Host | 1.700 | 0.01909 | **0.003** |
| Crop History | 1.634 | 0.00918 | **0.029** |
| Year ~ Compartment | 14.743 | 0.04139 | **0.001** |
| Year ~ Host | 2.548 | 0.02862 | **0.001** |
| Year ~ Crop History | 2.047 | 0.01149 | **0.005** |
| Compartment ~ Host | 1.257 | 0.01412 | 0.071 |
| Compartment ~ Crop History | 1.257 | 0.00706 | 0.139 |
| Host ~ Crop History | 0.993 | 0.02232 | 0.458 |
| Year ~ Compartment ~ Host | 1.931 | 0.02169 | **0.001** |
| Year ~ Compartment ~ Crop History | 1.325 | 0.00744 | 0.087 |

**Table S4**. Nitrogen (N), phosphorous (P), potassium (K), and sulfur (S), available in the soil, and fertiziler applied, at Swift Current, Saskatchewan, over each year of the two-year crop rotation experiments. Adapted from Hossain *et al*., 2019.

| | Swift Current | | |
|---|---|---|---|
| | 2015 | 2016 | 2017 |
| Available N-$P_2O_5$-$K_2O$-S (kg ha $^{-1}$)[a] | | | |
| Chem-fallow | 45-48-652-25 | 37-34-646-22 | 42-34-446-83 |
| Lentil | 24-47-728-31 | 20-33-578-28 | 30-43-488-83 |
| Wheat | 22-41-621-21 | 18-31-511-19 | 18-26-482-82 |
| Fertilizer Applied N-$P_2O_5$-$K_2O$-S (kg ha $^{-1}$) | | | |
| Chem-fallow | 45-7-0-10 | 48-7-0-10 | 55-7-0-10 |
| Lentil | 65-7-0-10 | 65-7-0-10 | 43-7-0-10 |
| Wheat | 65-7-0-10 | 68-7-0-10 | 67-7-0-10 |

a, Measurements taken at planting with soil available N and S at 0–60 cm depth, P and K at 0–15 cm depth.

#### Supplementary Figures

**Fig. S1**

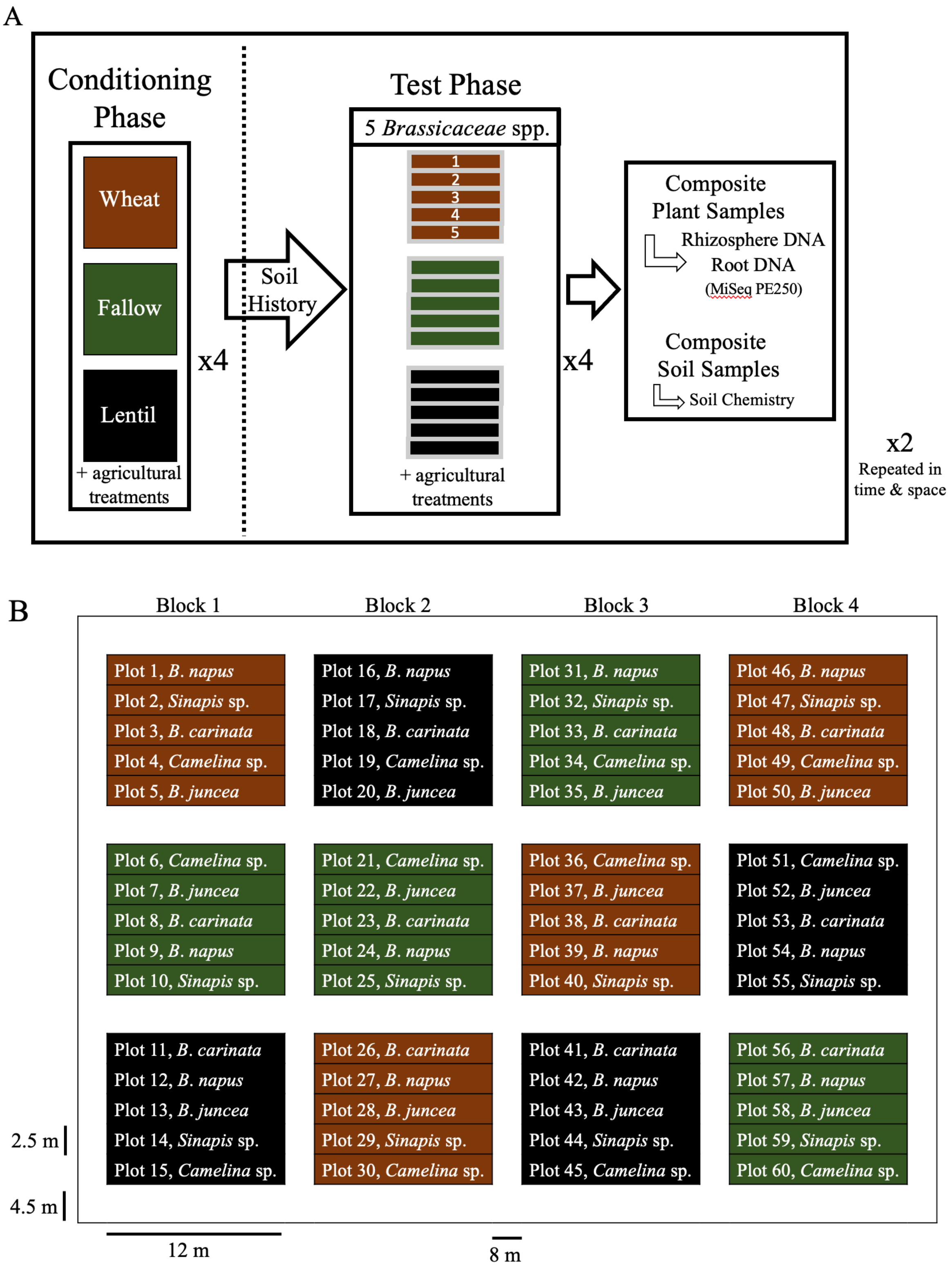

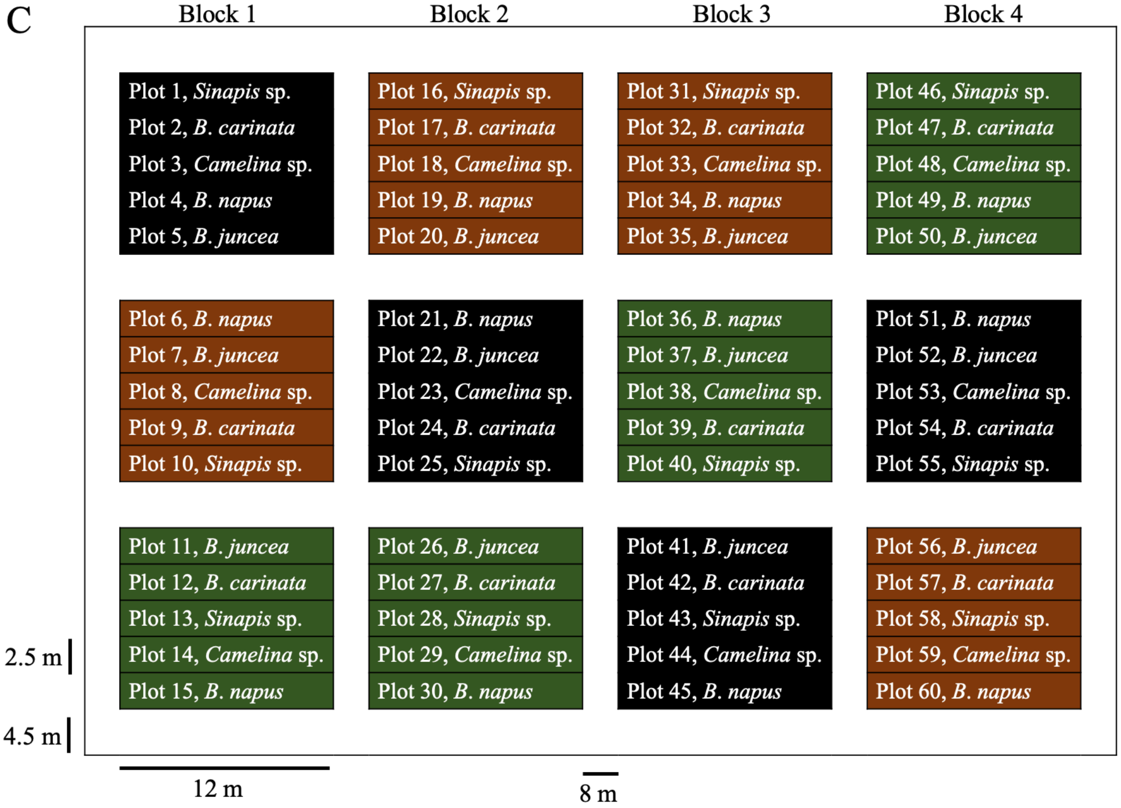


Figure S1. A two-year, or phase, cropping sequence was established in a field previously growing spring wheat (*Triticum aestivum* cultivar AC Lillian). The experimental design was a split-plot replicated in four complete blocks. In Phase 1, the 'Conditioning Phase', three soil history treatments were randomly assigned, consisting of spring wheat (*Triticum aestivum*, cv. AC Lillian), red lentil (*Lens culinaris* cv. CDC Maxim CL), or left fallow (brown, black, green, respectively). In Phase 2, the 'Test Phase', the conditioned plots were each subdivided and five *Brassicaceae* oilseed crop species were randomly assigned to one of these five subplots. Thus, each experiment had 60 subplots to sample. The experiment was performed twice, Experiment 1, 2015-2016, and Experiment 2, 2016-2017, on adjacent sites. (B) Experiment 1 field plan for the *Brassicaceae* crops, which were Ethiopian mustard (*Brassica carinata* L., cv. ACC110), canola (*B. napus* L., cv. L252LL), oriental mustard (*B. juncea* L., cv. Cutlass), yellow mustard (*Sinapis alba* L., cv. Andante), and camelia (*Camelina sativa* L., cv. Midas). Boarder space between plots and blocks is in white. (C) Experiment 2 field plan for the same *Brassicaceae* crops. For further details of this well-described experiment and its design, see Hossain *et al*. (2019), Liu *et al*. (2019), and Wang *et al*. (2020).

**Fig. S2**

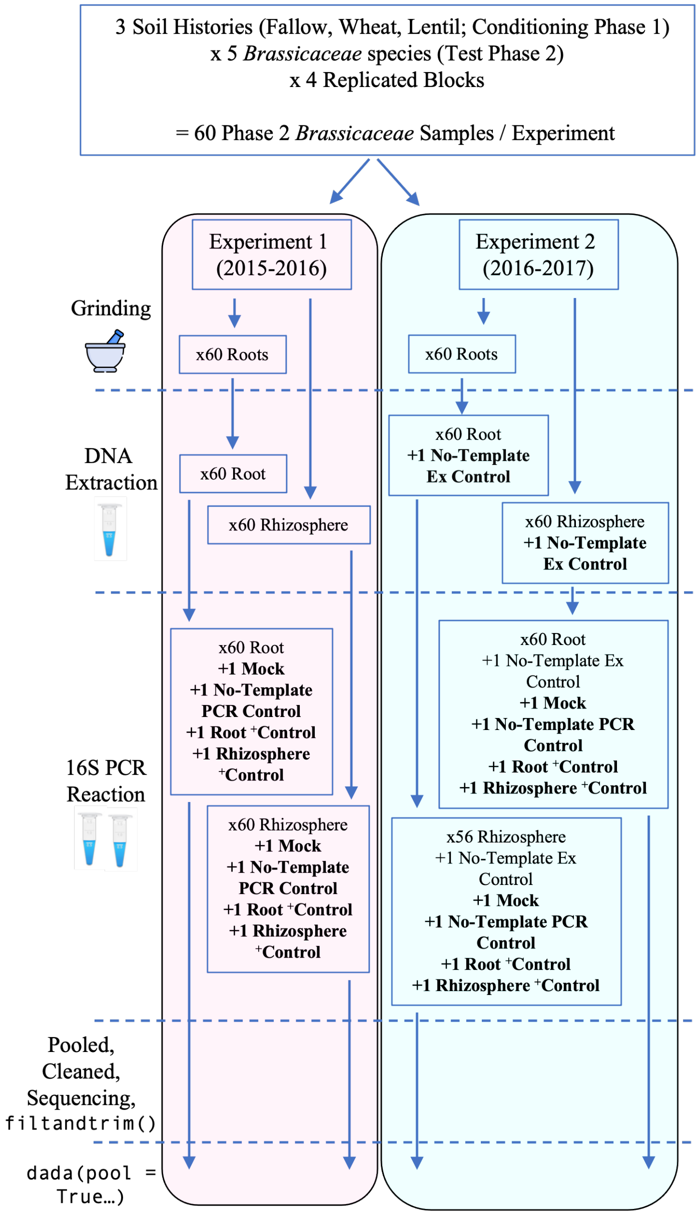


Figure S2. Organization of our lab workflow for the Test Phase *Brassicaceae* samples from harvest to generating amplicon sequence variants (ASVs). The Test Phase *Brassicaceae* samples were harvested in mid-late July. Four plants from two different locations within each of the 60 subplots were excavated and pooled together as a composite sample (Hossain *et al*., 2019; Liu *et al*., 2019, Wang *et al*., 2020). In the field, each plant had its rhizosphere soil divided from the root material, both portions were immediately flash-frozen in liquid nitrogen, and kept on ice. In the lab, roots were ground in liquid nitrogen, and DNA was extracted from all the Test Phase *Brassicaceae* root and rhizosphere portions. No-template extraction controls were included to assess what contaminates, or biases, the extraction kits might impart. All DNA samples were used as templates for PCR amplification of the 16S rRNA gene as a metabarcode. All the samples were PCR amplified twice, in two independent reactions, except four rhizosphere samples from experiment 2, which we were unable to be amplify, and were subsequently excluded. We included root and rhizosphere DNA from a previous experiment as PCR positive controls, as well as a bacterial mock to assess the accuracy, bias, and sensitivity of the lab workflow and bioinformatics pipeline. We also included no-template PCR negative controls, and confirmed by gel electrophoresis that none of the no-template extraction controls, nor the no-template PCR controls, contained DNA prior to sequencing. The two independent PCR reactions were pooled together for each sample, and all controls, cleaned and submitted for paired-end 250 bp Illumina MiSeq sequencing. To help identify sequencing biases, or bath effects, a replicate of the bacterial mock community was included on each of the four plates submitted for sequencing. All reads were subsequently trimmed and processed through the DADA2 pipeline for ASV inference.

Fig. S3

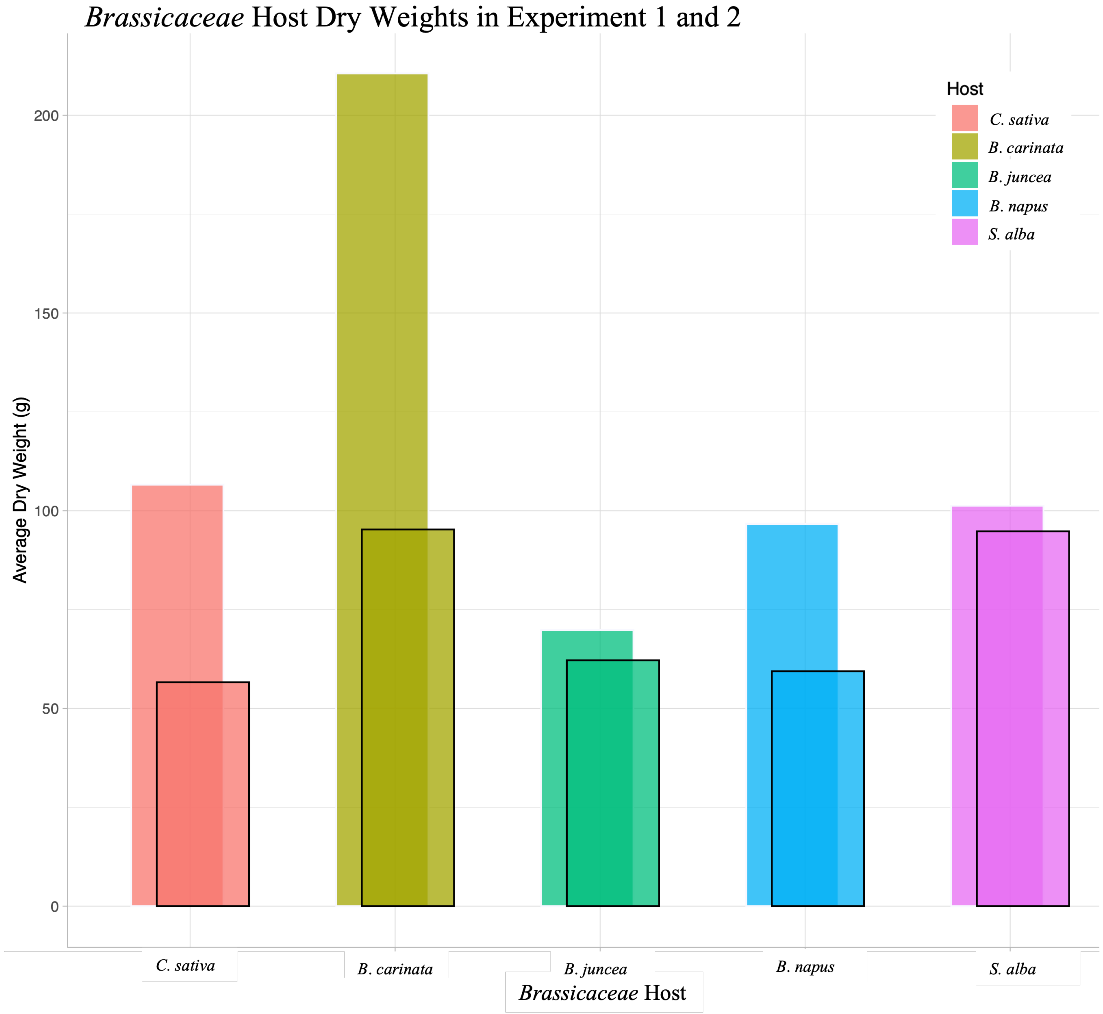


Figure S3. *Brassicaceae* host dry weights (g) decreased in Experiment 2 (black outlines), compared to Experiment 1 (no outlines). The Test Phase *Brassicaceae* samples were harvested in mid-late July, at Swift Current, Saskatchewan. The aerial portions were retained and dried to determine their weight.

Fig. S4

C

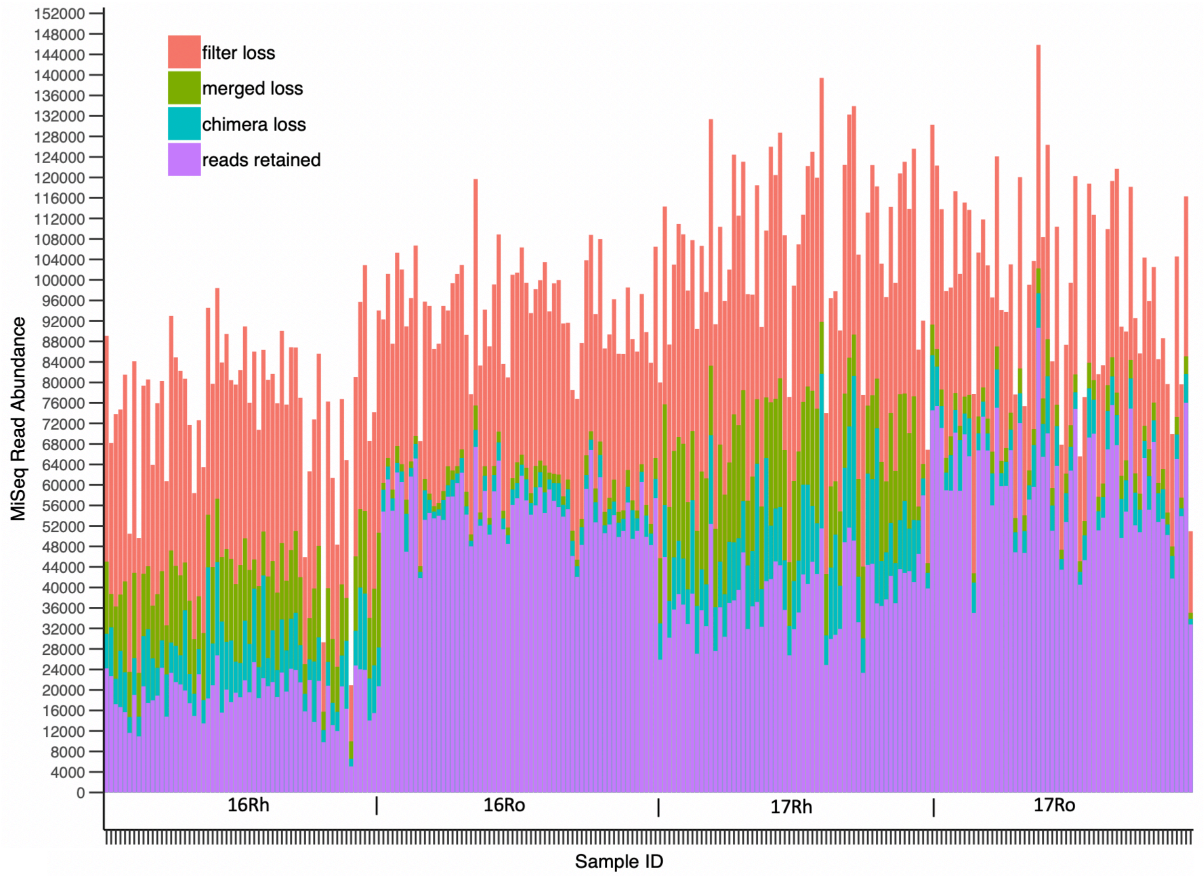


Figure S4. The DADA2 workflow processed 23 313 756 raw reads produced from one lane of sequencing via Illumina's MiSeq at Génome Québec in order to infer amplicon sequence variants (ASVs). A) Reads were produced in each of the six no-template negative controls, although no DNA was detected in these samples post-extraction and post-PCR. The stringency of the `filterAndTrim` step eliminated nearly all the reads from these samples. B) A replicate of the mock community was included on each plate submitted for sequencing: a rhizosphere and root plate for experiment 1, and a rhizosphere and root plate for Experiment 2. All four replicates of the bacterial mock community were very uniform in their processing, and yielded the recommended standard of approximately a third of the reads retained. C) An average of 43 129 ±18 032 high-quality reads per Test Phase *Brassicaceae* sample, for a total of 10 178 467, were retained. As observed in the no-template negative controls, and the bacterial mock community, prepared with the Test Phase from Experiment 2 samples, the Experiment 2 samples tended to have more reads than the Experiment 1 samples. Among the Test Phase Brassicaceae samples, the root samples retained noticeably more reads than their rhizosphere partners. Note that root samples also retained more reads during the merging and chimera steps in DADA2 than the rhizosphere samples. There was no evident difference in reads among the five *Brassicaceae* species, beyond the effects of compartment and year.

Fig. S5

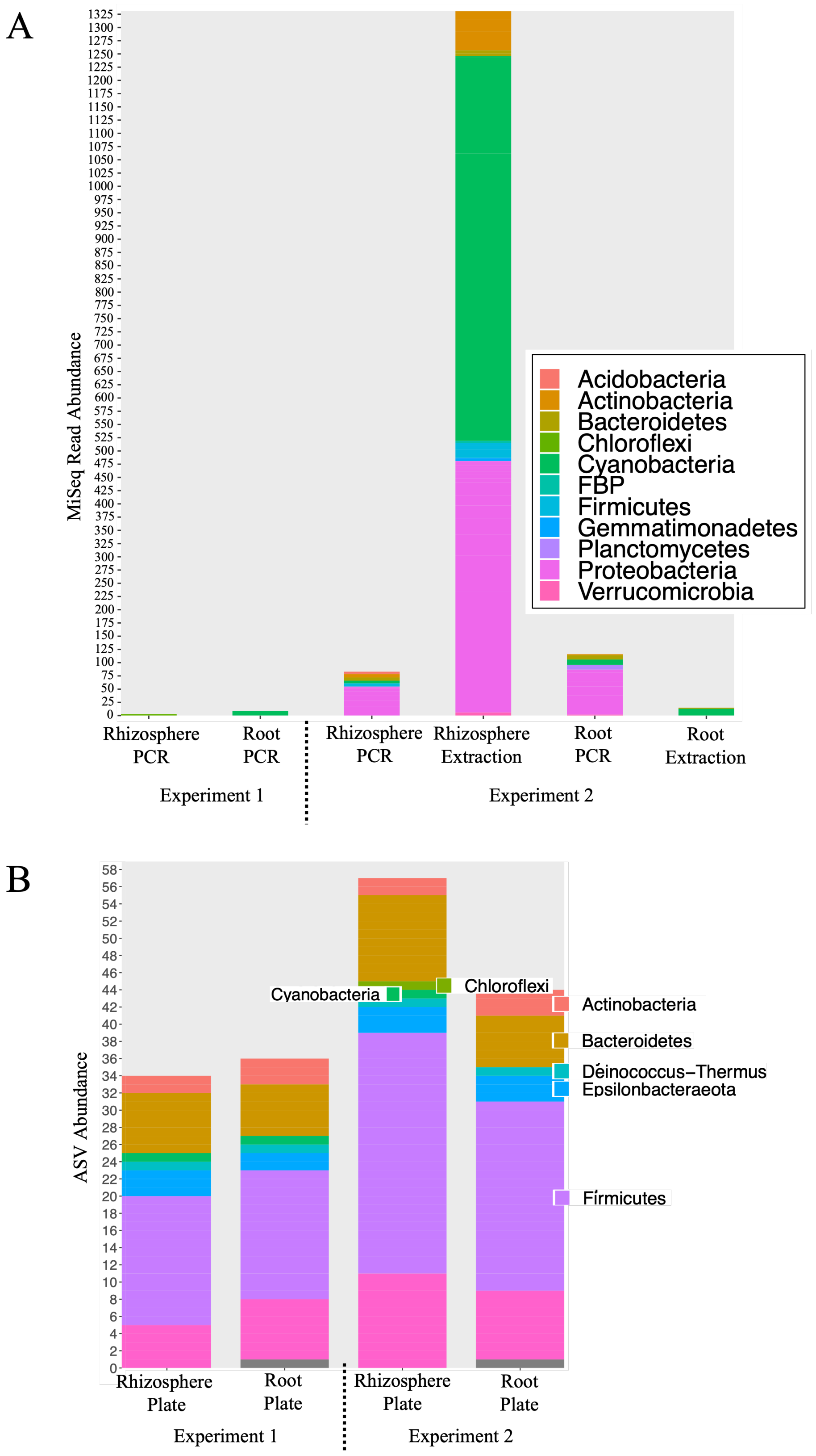
A
MiSeq Read Abundance
Acidobacteria
Actinobacteria
Bacteroidetes
Chloroflexi
Cyanobacteria
FBP
Firmicutes
Gemmatimonadetes
Planctomycetes
Proteobacteria
Verrucomicrobia
Rhizosphere PCR
Root PCR
Rhizosphere PCR
Rhizosphere Extraction
Root PCR
Root Extraction
Experiment 1
Experiment 2
B
ASV Abundance
Cyanobacteria
Chloroflexi
Actinobacteria
Bacteroidetes
Deinococcus−Thermus
Epsilonbacteraeota
Firmicutes
Rhizosphere Plate
Root Plate
Rhizosphere Plate
Root Plate
Experiment 1
Experiment 2

Figure S5. High-quality MiSeq reads retained through the DADA2 pipeline from among the no-template negative controls, and mock community replicates, were inferred as inferred amplicon sequence variants (ASVs), and assigned taxonomy using the Silva database, represented here as phlya. A) ASVs inferred from among the six no-template negative controls, represented as their corresponding abundance of reads to illustrate these ASVs were derived from few reads. The negative controls had between 1 and 32 different ASVs, where most reads that were retained were inferred as ASVs identified as *Cyanobacteria* and *Proteobacteria*. B) 34 – 55 individual ASVs were inferred from among the four mock community replicates; the mock replicate prepared with the experiment 2 rhizosphere samples, followed by the mock replicate in the experiment 2 root sample, contained the most ASVs. Every bacterial group included in the mock community was detected in each replicate. The *Actinobacteria*, and *Deinococcus*, were the most specific, as 2, or 1, species were included in the mock community, respectively, and accurately detected in our bioinformatics pipeline. There was an expansion among the ASVs identified as *Bacteriodetes*, *Firmicutes*, and *Proteobacteria*. Three of the mock replicates had an ASV identified as a *Cyanobacteria*, while one of them also replicates had an ASV identified as a *Chloroflexi*, though none were included in the community's composition.

Fig. S6

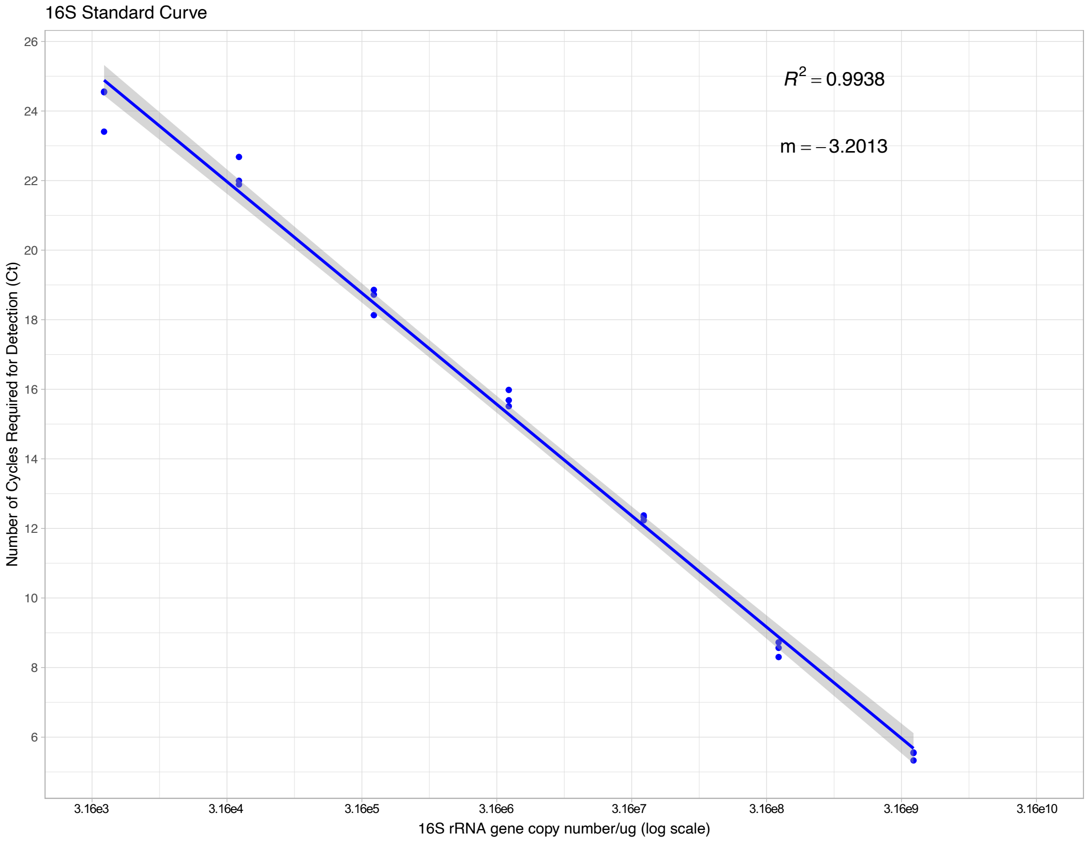


Figure S6. A standard curve of the 16S rRNA gene copy numbers (X-axis) versus the number of cycles required for detection (Ct, Y-axis), as determined from the serial dilution of a quantified 16S rRNA gene.

Fig. S7A

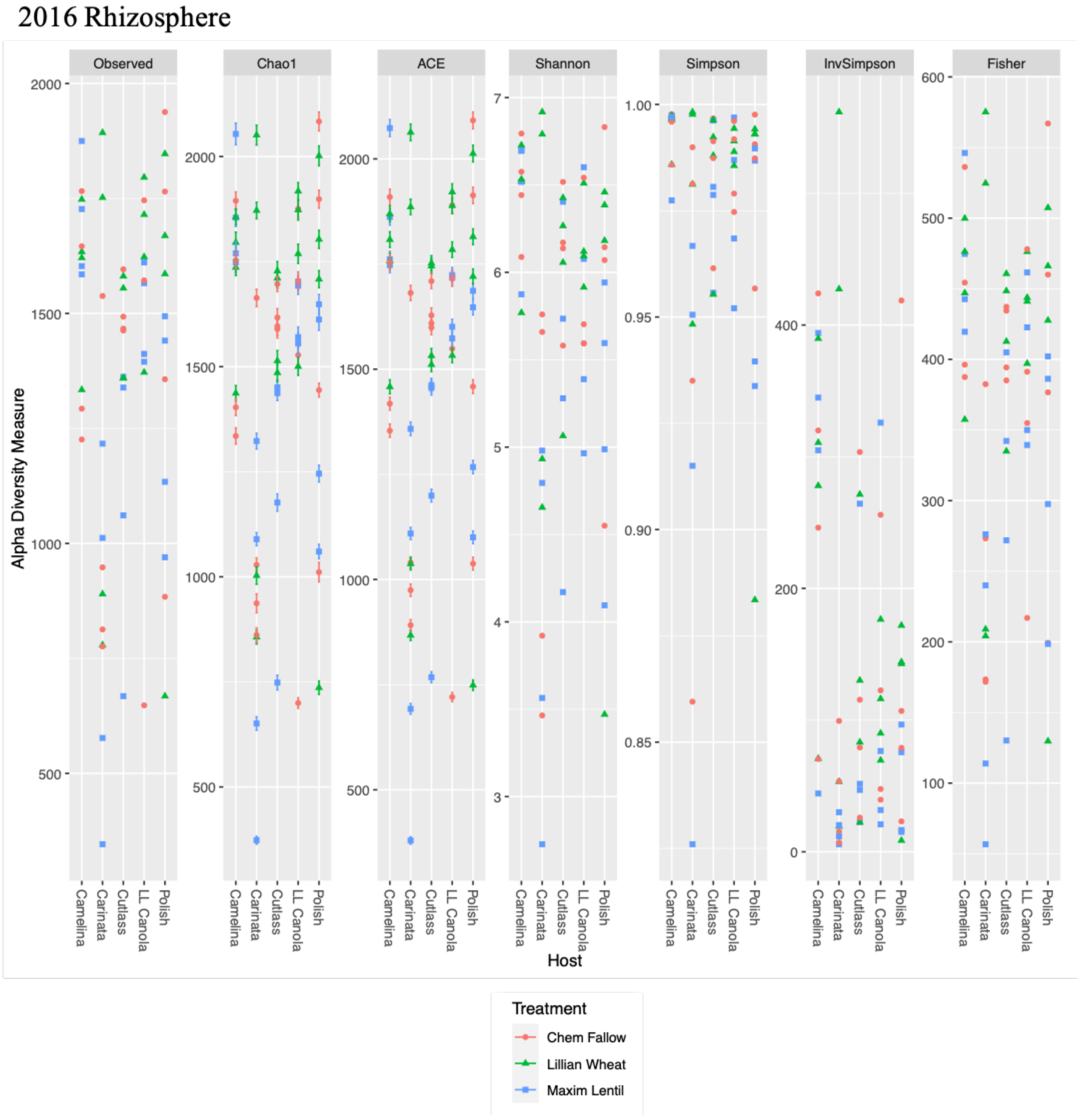
2016 Rhizosphere
Observed
Chao1
ACE
Shannon
Simpson
InvSimpson
Fisher
Alpha Diversity Measure
Host
Camelina
Carinata
Cutlass
LL Canola
Polish
Treatment
Chem Fallow
Lillian Wheat
Maxim Lentil

Fig. S7B

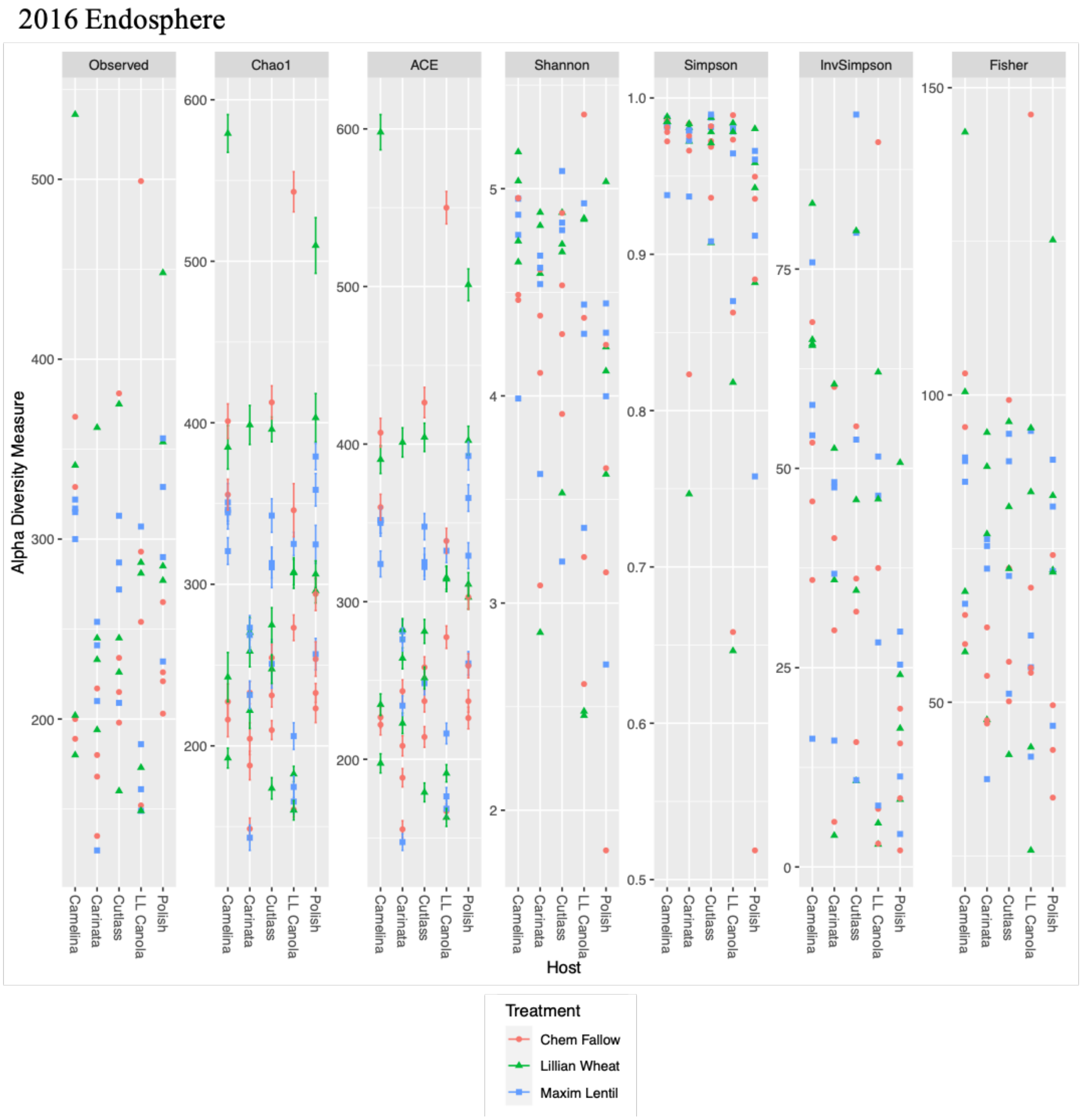


Figure S7. Taxa-based α-diversity indices (y-axis) for the rhizosphere (A) and root (B) communities from Experiment 1, harvested 2016. Each α-diversity index was grouped by *Brassicaceae* host, and reflect the phylogenetic diversity observed, where communities are broadly similar across hosts, and soil histories. Similar results were also observed for the communities based on relative abundance, as well as in 2017 (data not shown).

Fig S8A

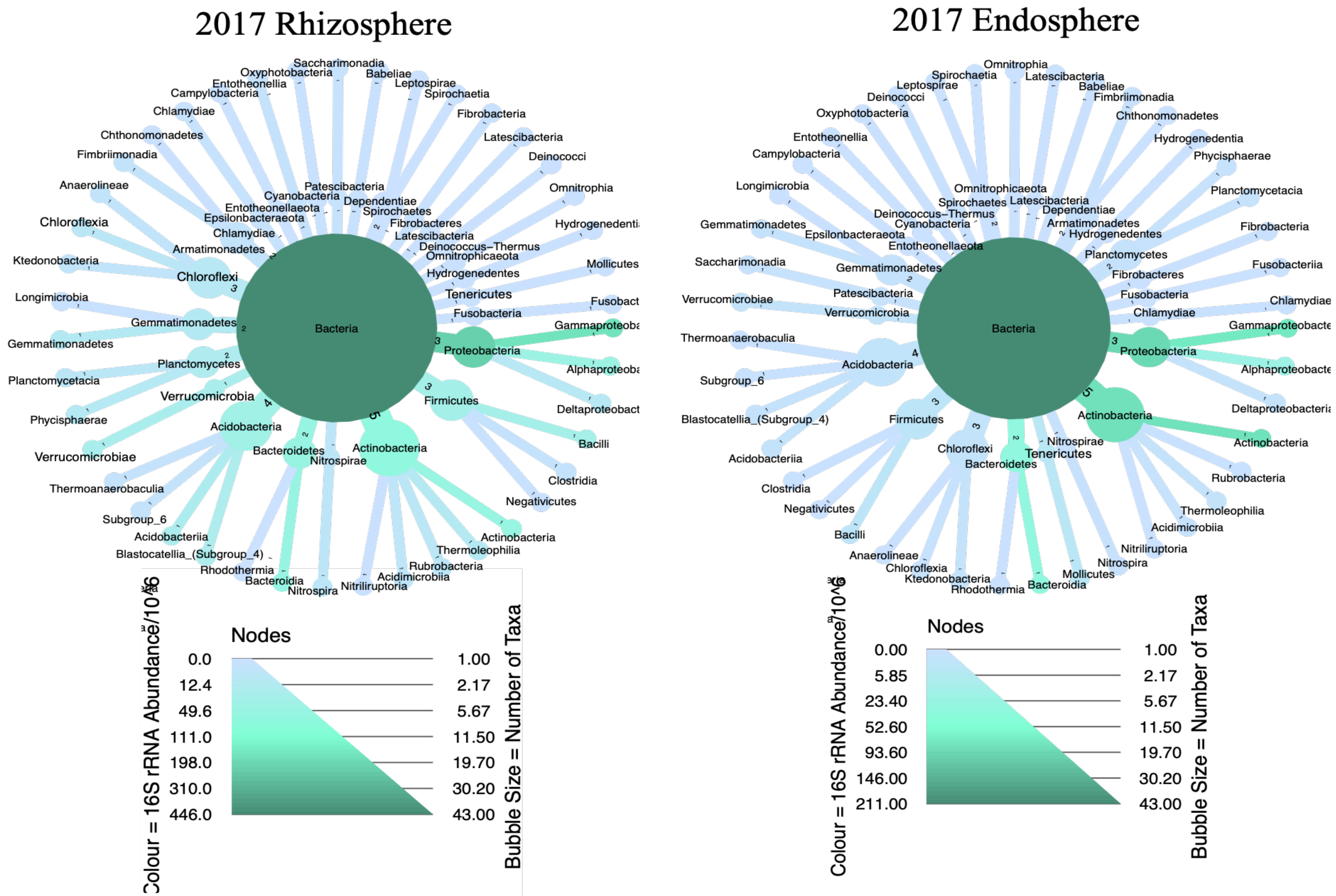

2017 Rhizosphere
2017 Endosphere
Bacteria
Proteobacteria
Actinobacteria
Acidobacteria
Firmicutes
Chloroflexi
Bacteroidetes
Nodes
Colour = 16S rRNA Abundance/10^6
Bubble Size = Number of Taxa
0.0
12.4
49.6
111.0
198.0
310.0
446.0
1.00
2.17
5.67
11.50
19.70
30.20
43.00
0.00
5.85
23.40
52.60
93.60
146.00
211.00

Fig. S8B

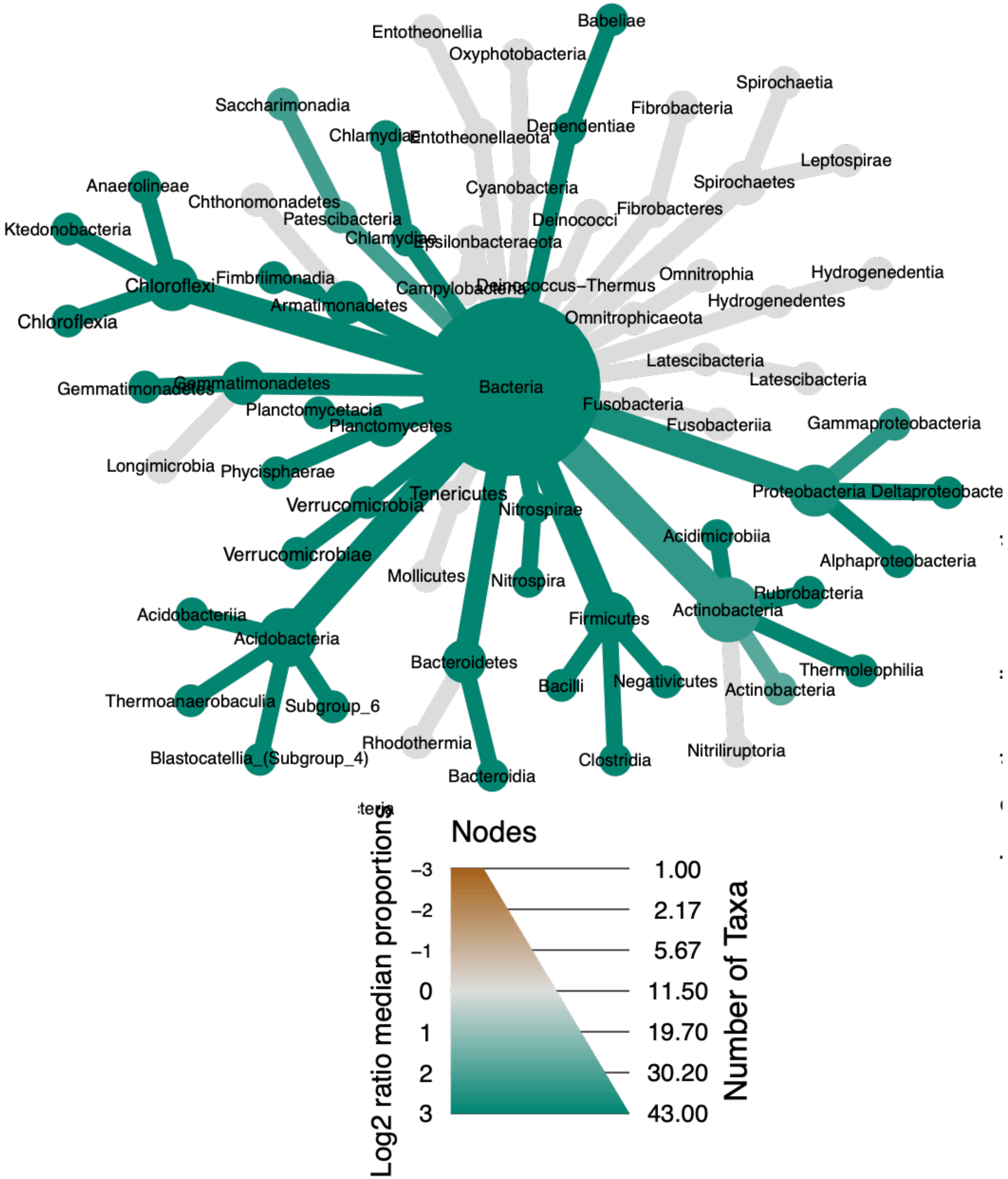
Bacteria
Babeliae
Entotheonellia
Oxyphotobacteria
Spirochaetia
Saccharimonadia
Fibrobacteria
Dependentiae
Chlamydiae
Entotheonellaeota
Leptospirae
Anaerolineae
Cyanobacteria
Spirochaetes
Chthonomonadetes
Patescibacteria
Deinococci
Fibrobacteres
Ktedonobacteria
Chlamydiae
Epsilonbacteraeota
Fimbriimonadia
Omnitrophia
Hydrogenedentia
Chloroflexi
Campylobacteria
Deinococcus−Thermus
Armatimonadetes
Hydrogenedentes
Chloroflexia
Omnitrophicaeota
Latescibacteria
Latescibacteria
Gemmatimonadetes
Gemmatimonadetes
Fusobacteria
Planctomycetacia
Fusobacteriia
Gammaproteobacteria
Planctomycetes
Longimicrobia
Phycisphaerae
Proteobacteria
Deltaproteobacte
Tenericutes
Verrucomicrobia
Nitrospirae
Acidimicrobiia
Verrucomicrobiae
Alphaproteobacteria
Mollicutes
Nitrospira
Rubrobacteria
Acidobacteriia
Actinobacteria
Firmicutes
Acidobacteria
Bacteroidetes
Thermoleophilia
Bacilli
Negativicutes
Actinobacteria
Thermoanaerobaculia
Subgroup_6
Rhodothermia
Nitriliruptoria
Blastocatellia_(Subgroup_4)
Clostridia
Bacteroidia
Nodes
Log2 ratio median proportions
−3
−2
−1
0
1
2
3
1.00
2.17
5.67
11.50
19.70
30.20
43.00
Number of Taxa

Figure S8. (A) Taxa clusters of the ASVs inferred from among the rhizosphere (left) and root (right) of the Test Phase bacterial communities from Experiment 2, harvested 2017, represented to the class level. The size of the taxonomic groups (bubbles) represents the number of ASVs that occured, and the colour scale represents the absolute abundance of each ASV. (B) The differential taxa cluster between the absolute abundance of the rhizosphere and root in Experiment 1, where the abundance of each taxonomic group in the cluster is compared between each compartment, using the using the non-parametric Kruskal test and the post-hoc pairwise Wilcox test, with the FDR correction. Taxa that are significantly (*p*. adj < 0.05) more abundant in rhizosphere are highlighted in green.

Fig S9

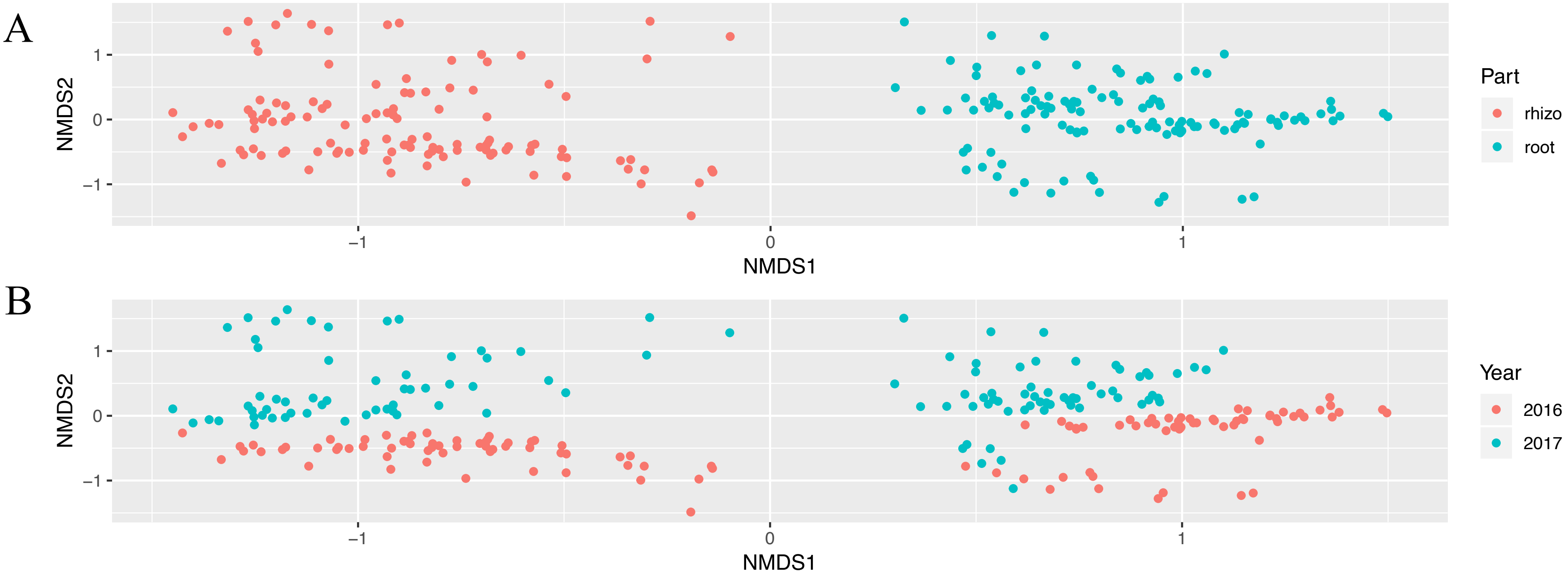


Figure S9. A non-metric multidimensional scaling (NMDS) of all the Test Phase data illustrates that the most important factors shaping the bacterial communities are: A) compartment, rhizosphere (orange), or root (turqoise) (PERMANOVA, $R^2 = 0.19304$, $p = 0.001$), and B) sampling year, 2016 (Experiment 1, orange), or 2017 (Experiment 2, turqoise) (PERMANOVA, $R^2 = 0.07067$, $p = 0.001$). As all the bacterial communities appear clustered together respectively by compartment and year, regardless of other factors, we can be confident that the data is not excessively biased, and batch effects have been subsumed within these larger factors/patterns. See Table S3 for these PERMANOVA results.

Fig. S10

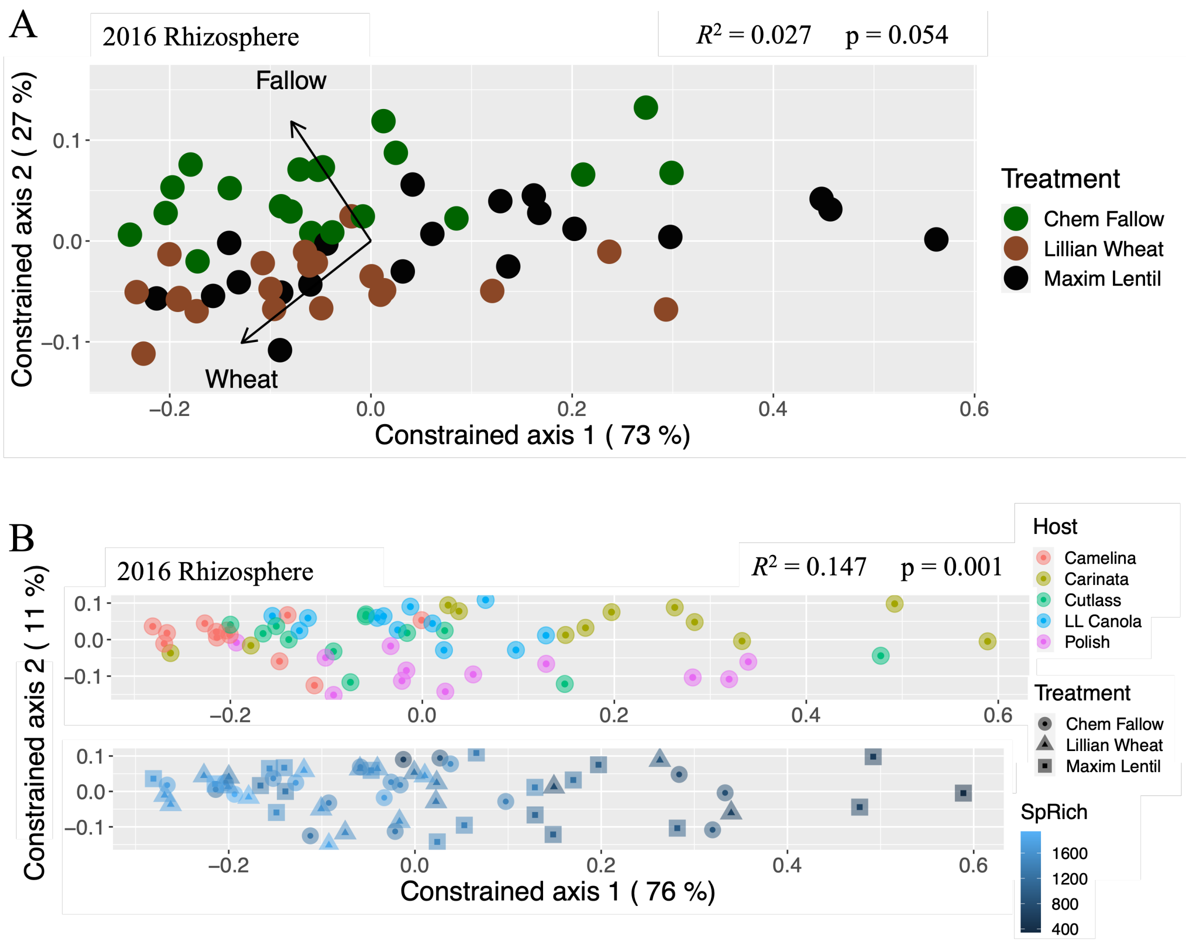


Figure S10. Bacterial rhizosphere communities exhibited significant structure according to (A) soil history, and (B) *Brassicaceae* host plants when harvested in 2016 from the Test Phase in Experiment 1 of a two-year crop rotation, in Swift Current, Sask. Weighted UniFrac distances were used with a distance-based redundancy analysis. (A) Soil histories established in the Conditioning Phase were significant ($R^2 = 0.027$, $p = 0.054$) a year later in structuring the bacterial communities from the Test Phase rhizosphere communities in Experiment 1, as illustrated by phylogenetically similar communities being more distinctive according to their soil histories. (B) *C. sativa* (red) bacterial rhizosphere communities were more phylogenetically similar compared to communities from other *Brassicaceae* hosts ($R^2 = 0.147$, $p = 0.001$). Communities from *Brassica carinata* appeared the least structured by host, and strongly impacted by lower species richness (SpRich). Note that the Test Phase bacterial rhizosphere communities in Experiment 1 were not significantly structured by soil chemistry (data not shown).

Fig. S11

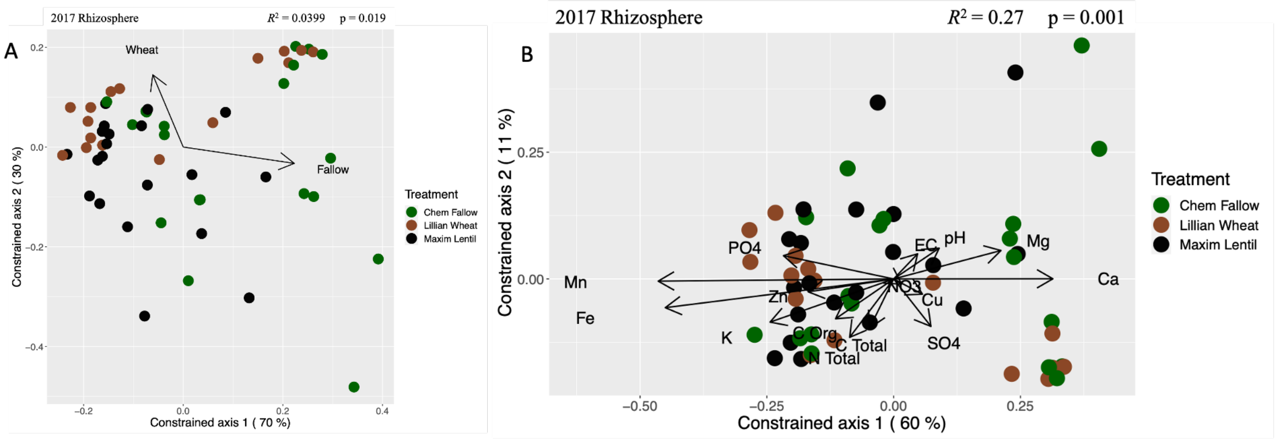


Figure S11. Bacterial rhizosphere communities exhibited significant structure according to (A) soil history, and (B) soil chemistry in Experiment 2, harvested in 2017, from Test Phase of a two-year crop rotation, in Swift Current, Sask. Weighted UniFrac distances were used with a distance-based redundancy analysis. (A) Soil histories established in the Conditioning Phase were still significant a year later ($R^2$ = 0.0399, $p$ = 0.019) in structuring the bacterial communities from the Test Phase rhizosphere communities in Experiment 2, though it is difficult to observe a clear trend among phylogenetically similar communities being more distinctive according to their soil histories. (B) Soil chemistry was also significant in structuring the Test Phase bacterial rhizosphere communities ($R^2$ = 0.27, $p$ = 0.001): manganese was contrasted by calcium, iron and zinc contrasted with magnesium, while potassium was opposed by pH, though there is not a clear trend among phylogenetically similar communities being more distinctive according to their soil chemistries. Note that the Test Phase bacterial rhizosphere communities from Experiment 2 were not significantly structured by *Brassicaceae* host plants (data not shown).

Fig. S12

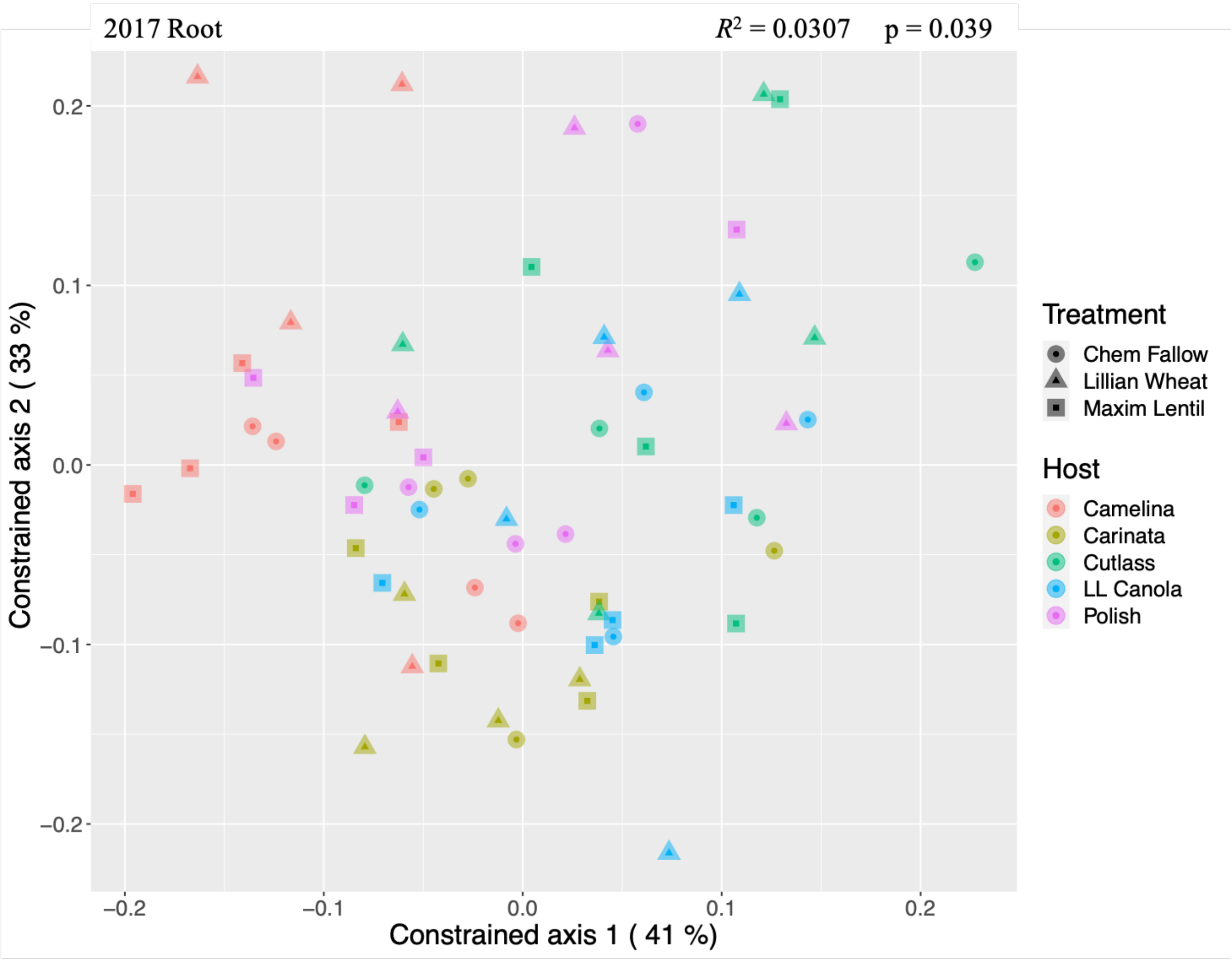


Figure S12. Bacterial root communities from Experiment 2 exhibited significant structure according to their *Brassicaceae* host plants when harvested in 2017 from the Test Phase of a two-year crop rotation, in Swift Current, Sask. Weighted UniFrac distances were used with a distance-based redundancy analysis. *Brassicaceae* host plants were significant ($R^2 = 0.0307$, $p = 0.039$) in structuring the Test Phase bacterial root communities. We observed a gradient between *C. sativa* (red, upper left), *Brassica juncea* cv. Cutlass (green, upper right), and *B. carinata* (brown, middle).
